# Variations of autonomic arousal mediate the reportability of mind-blanking occurrences

**DOI:** 10.1101/2024.03.26.586648

**Authors:** Boulakis Paradeisios Alexandros, Simos Nicholas John, Zoi Stefania, Mortaheb Sepehr, Schmidt Christina, Raimondo Federico, Demertzi Athena

## Abstract

Mind-blanking (MB) is the inability to report mental events during unconstraint thinking. Previous work shows that MB is linked to decreased levels of cortical arousal, indicating dominance of cerebral mechanisms when reporting mental states. What remains inconclusive is whether MB can also ensue from autonomic arousal manipulations, pointing to the implication of peripheral physiology to mental events. Using experience-sampling, neural, and physiological measurements in 26 participants, we first show that MB was reported more frequently in low arousal conditions, elicited by sleep deprivation. Also, there was partial evidence for a higher number of MB reports in high arousal conditions, elicited by intense physical exercise. Transition probabilities revealed that, after sleep deprivation, mind-wandering was more likely to be followed by MB and less likely to be followed by more mind-wandering reports. Using classification schemes, we show higher performance of a balanced random forest classifier trained on both neural and physiological markers in comparison to performance when solely neural or physiological were used. Collectively, we show that both cortical and autonomic arousal affect MB report occurrences. Our results establish that MB is supported by combined brain-body configurations, and, by linking mental and physiological states they pave the way for novel, embodied accounts of spontaneous thinking.

## Introduction

During ongoing mentation, our mind constantly shifts across different mental states. These mental states typically bear some content (“what we think about”) and indicate a relationship towards that content (i.e., perceiving, fearing, hoping, remembering) [1]. As we move through the environment, our thoughts fluctuate between the external and internal milieu [2, 3], resulting in a fluid stream of consciousness [4]. External content is tightly coupled to the processing of environmental stimuli and task-demanding conditions. Internal content is more associated with self-referential processing and internal dialogue, widely known as Mind-Wandering (MW) [4]. Inclusive as this external-internal dipole may seem, it does not capture the full scope of the “aboutness” of mental content. Recent work has highlighted another mental state, where people report that they are “thinking of nothing” or “their mind just went away”, a phenomenological experience termed mind-blanking (MB) [5]. As MB is relatively new in the landscape of ongoing cognition, the extent of MB episodes in daily and clinical settings remains widely uncharacterized. For example, a recent study found that MB might be miscategorized as MW in ADHD symptom evaluation [6]. Therefore, the experience of MB occurrences poses a challenge to our everyday functioning and our understanding of the continuous nature of the stream of consciousness.

Currently, there is no clear answer as to how MB reports are generated. So far, behavioral studies show that MB can arise after conscious mental effort to empty our mind [7, 8, 9], is usually unintentional [5, 10, 11] and gets reported less frequently during unconstrained thinking compared to MW and sensory/perceptual mental states [5, 11, 12, 13]. At the brain level, the inability to report mental events after the prompt to “empty the mind” has been associated with activation of the anterior cingulate/medial prefrontal cortex, and deactivation of inferior frontal gyrus/Broca’s areas and the hippocampus, which the authors interpreted as the inability to verbalize internal mentation (inner speech) [8]. Recently, we found that the functional connectome of fMRI volumes around MB reports was similar to a unique brain pattern of overall positive inter-areal connectivity [12] which was also characterized by increased amplitude of fMRI global signal (i.e. averaged connectivity across all grey matter voxels), an implicit indicator of low arousal [14, 15, 16]. For example, the amplitude of the global signal correlated negatively with EEG vigilance markers (alpha, beta oscillations), while increases in EEG vigilance due to caffeine ingestion were associated with reduced global signal amplitude [14]. Our findings corroborate recent EEG-related evidence supporting the possibility of “local sleeps” during MB reportability [10, 17]. “Local sleeps” refer to the scalp distribution of EEG potentials during wakefulness, in the form of high-intensity, slow oscillatory activity in the theta/delta band, which could differentiate between MB and MW, with more frontocentral potentials tied to MW and parietal to MB [10]. Together, the presence of slow waves preceding MB reports and the high fMRI global signal hint toward the role of arousal in mental content reportability. Starting from this line of evidence, we generally infer that arousal fluctuations drive MB reportability.

Arousal is a multidimensional term generally referring to the behavioral state of being awake and alert, supporting wakefulness, responsiveness to environmental stimuli, and attentiveness [18, 19]. Anatomically, arousal is supported by the ascending arousal system, the autonomic nervous system, and the endocrine system [18]. Early on, Lacey viewed arousal in terms of behavioral arousal (indicated by a responding organism, like restlessness and crying), cortical arousal (evidenced by desynchronized fast oscillatory activity), and autonomic arousal (indicated by changes in bodily functions) [20]. Cortical arousal is self-generated through the reticulate formation and propagated through dorsal, thalamic, and ventral subthalamic pathways [21], and can be indexed by the alpha, theta, and delta EEG bands during wakefulness [22, 23]. Lower levels of cortical arousal in the form of slow waves have been associated with an increased number of missed stimuli in behavioral tasks [11, 23] and decreased thought intensity [24]. Also, lower levels of arousal indexed by pupil size have been correlated with a higher probability of MB reports in sustained attention tasks [11, 25, 26].

Much as it may have been done in terms of cortical arousal, the present study will focus on how autonomic arousal influences MB reportability, which is widely understudied. Our choice is justified by the theoretical assumption that mental function is tightly linked to peripheral body functions, explicitly expressed by the embodied cognition stance [27]. Briefly, embodiment holds that cognition is bound to a living body interacting with a dynamic environment and conceptualizes cognition as the result of brain-body interactions during dynamic contexts. From that perspective, modifications in autonomic arousal are expected to lead to differential reportability of mental states. Autonomic arousal links the body and the brain through spinal cord projections from peripheral organs to the brainstem and can be indexed by physiological signals reflecting sympathetic/parasympathetic balance, such as heart rate, galvanic skin response, and fluctuations in pupil size [28]. Converging evidence suggests that afferent physiological signals and biological rhythms, such as the cardiac or the respiratory phase, play a modulatory role in conscious perception [29, 30], metacognition [31], affective salience of information [32], and perceptual confidence of sensory sampling [33], both during task performance and in-silico simulations [34]. Alterations in autonomic arousal were also found to influence brain activity in that fMRI volumes characterized by lower arousal levels (indexed by decreased pupil size), showed reduced in-between network integration and inter-subject variability in comparison to scans characterized by high arousal levels (indexed by increased pupil size) [35].

Taken together, we here advocate for a direct link between autonomic arousal and content reportability. Firstly, we examined how MB report distribution shifted across different autonomic arousal conditions. To this end, we used experience-sampling under differently elicited arousal conditions. Experience-sampling is a though-sampling methodology, where people are probed to report their mental state at random intervals, probed by an external cue [4]. We employed this task at three distinct arousal conditions: *Baseline*, *High* (post-workout), and *Low* (post-sleep deprivation). Our operational hypothesis was that optimal levels of autonomic arousal (fixed variable) are necessary for optimal mental state reportability (dependent variable). We expected that deviations from optimal levels, such as after sleep deprivation or intense physical exercise, would alter our stream of thought, therefore promoting more frequent MB reports (Supplementary Table S1 for the full scope of our hypothesis). Secondly, we opted to identify specific brain-body interaction patterns that would promote MB reportability. To this end, we utilized multimodal neurophysiological recordings and a machine-learning approach to decode MB reports from arousal measurements.

## Methods

### Ethics Information

The experimental procedure has been approved by the CHU Liège local ethics committee and conforms with the Declaration of Helsinki and the European General Data Protection Regulation (GDPR). Before the onset of the protocol, participants provided informed consent for their participation in the study. Participants also received monetary compensation for their participation in the study.

### Design

The study included healthy volunteers recruited after campus poster advertisements, intranet electronic invitations, and through the ULiège “petites annonces” e-campus platform. Inclusion criteria were: a) right-handedness, b) age>18 years, c) minimal exercise background (<2h per week), d) good subjective sleep quality (Pittsburgh Sleep Quality Index [PSQI] ≤ 5 [36]), e) habitual sleep duration of 8 ± 1 hours. Exclusion criteria were: a) history of developmental, psychiatric, or neurological illness resulting in documented functional disability, b) severe anomalies in pupil shape or inability to open both eyes preventing pupil measurement [37], c) analgesic medication which may affect physiological arousal, d) history of psychiatric illness pertaining to anxiety disorders or scores < 9 in the General Anxiety Disorder-7 (GAD-7 scale) [38] as anxious participants experience biased perceptions of their bodily states [39], e) extreme chronotypes, f) shift work or traveling over time zones in the past 3 months.

Experience-sampling was utilized in a within-participants repeated-measures design. During the experience-sampling session, participants laid restfully and were directed to let their minds wander, without any specific instructions towards internal (daydreaming, memories, prospective events) or external thoughts (body sensations, sensory stimuli in their immediate environment). Auditory probes (total n=40, 500Hz simple tones) invited participants to report what they were thinking at the moment just preceding the probe. The inter-probe interval was sampled from a uniform distribution between 110 and 120 seconds. Report times were monitored online to examine if participants missed the probe or fell asleep due to our experimental manipulation. In case of a report time > 6s, participants were reminded to report their mental state as soon as they heard the probe and indicate they were awake via button press. In case of unresponsiveness, the experimenters manually awakened the participant. Depending on the probes’ trigger times and participants’ reaction times, a recording lasted on average 70-90 minutes. We chose to present 40 probes (overall length approximately 1h and 15min) to avoid fatigue/drowsiness and the possibility of participants returning to baseline arousal after the experimental manipulations. Also, the relatively large experience-sampling interval, compared to previous studies, was used to record enough samples to accurately estimate physiological markers from slow oscillatory signals, such as heart-rate variability. Upon the probe, participants had to choose among four distinct choices describing their mental state: mind-blanking (MB), mind-wandering (MW), perceptual sensations (SENS), or sleep (SLEEP). These response options were chosen to minimize assumptions about what the actual partitions of mental states might be. For example, debates about what can be classified as MW [40] refer to whether MW is a coherent cluster of events [1, 41] and how it is separated from awareness and processing of environmental stimuli [40, 42]. We believe that our divisions respect the literature on internal/external thinking networks [3, 43, 44] while introducing minimum assumptions as to the actual content of each state. The introduction of the sleep option facilitated the identification of trials where participants fell asleep due to the experimental manipulation. Participants indicated their responses via button press from a response keyboard placed under their dominant hand. We repeated the experience-sampling task on three distinct days, over the span of two weeks under three conditions: a) experience-sampling under spontaneous thinking without arousal modulations (*Baseline*), b) experience-sampling elicited through short, high-intensity interval training (*High Arousal*), c) experience-sampling after total sleep deprivation (*Low Arousal*) (Fig. 1). The goal of both arousal manipulations was to promote distinct changes in physiological and cortical markers associated with arousal mechanisms (Supplementary Table S2). Monitoring of arousal changes was done with physiological and cortical measurements. In case when participants did not show distinct cortical and physiological changes after our arousal manipulations, they were excluded from further analysis. Effect monitoring was done by examining the heart rate in *High Arousal* as well as the EEG spectra in both *High* and *Low Arousal*.

**Figure 1.**
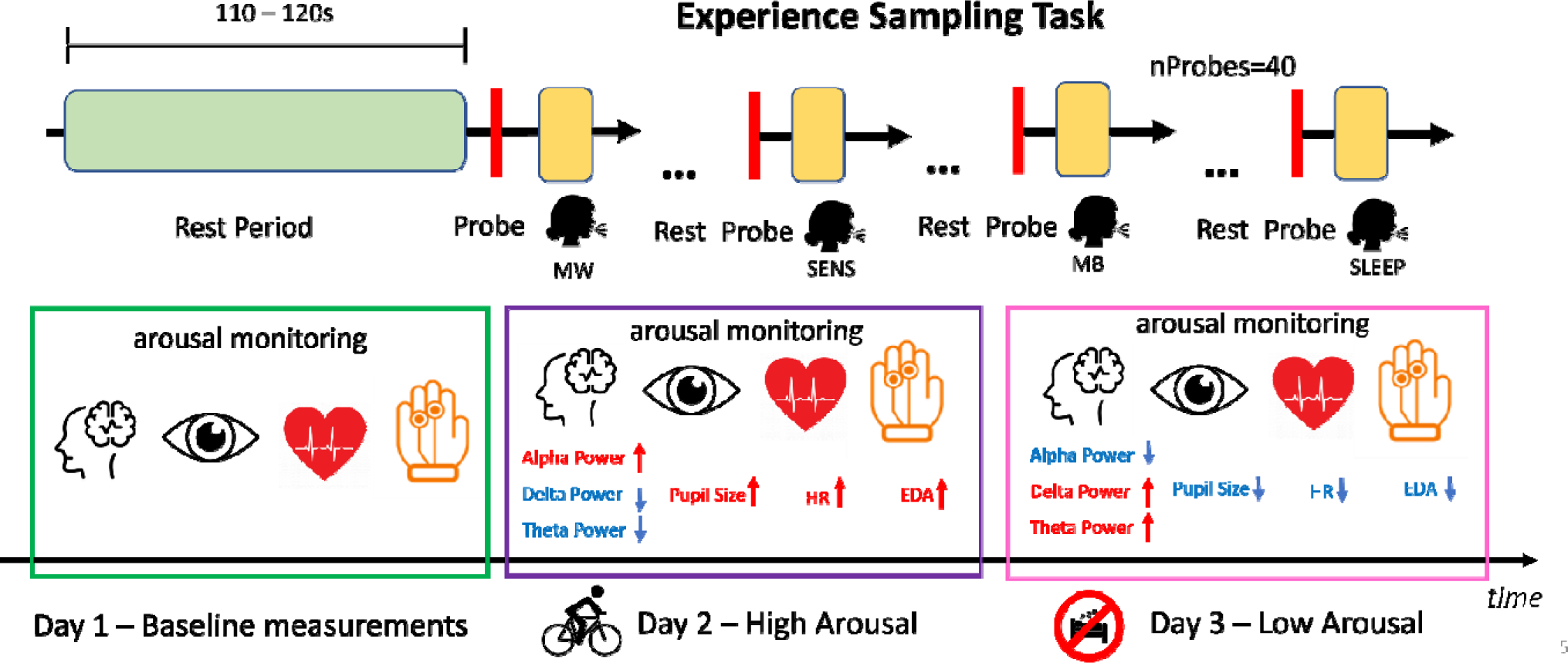
Experimental protocol. *Top* The experience-sampling task invited participants to sit idly and relax, letting their minds wander. Every 110-120s, a 500 Hz auditory cue probed participants to report what they were thinking at that moment. Participants were able to choose from 4 presented responses: Mind-Blanking (MB), Mind-wandering (MW), Perceptual Sensations (SENS), and Sleep (SLEEP). *Bottom* Repeated-measures autonomic arousal recordings. To test how spontaneous thoughts unfold over time across different arousal conditions, we first invited people for baseline assessments on Day 1 (*Baseline* condition). On Day 2 participants underwent a 15-minute high-intensity exercise routine (*High Arousal* condition) and on Day 3 they participated in a total sleep deprivation protocol (*Low Arousal* condition). The *High* and *Low Arousal* conditions were counter-balanced across participants. Multimodal physiological recordings were used to monitor arousal manipulations. The dataset was constituted of EEG, pupillometry, ECG, EDA, and respiratory data; the arrows indicate the hypothesized directions of the derived metrics.

In *High Arousal*, participants first performed high-intensity interval activity in the form of cycling. They started with a warm-up training session of 3 minutes to avoid potential muscle trauma and then cycled for 45 seconds as fast as possible. A resting period of 15 seconds followed. A total number of 10 workout cycles was administered. The choice of this timing protocol rested on previous studies indicating that similar exercise routines produce distinct and sustained sympathetic activity [45, 46] and cortical excitation [46], which can last between 30-90 minutes after exercise cessation[47].

In *Low Arousal*, participants performed the experience-sampling task after one night of total sleep deprivation. Sleep deprivation leads to an arousal state that is behaviourally distinct from typical wakefulness [48, 49], promotes specific neuronal signatures (“local sleeps” in the delta band) [11], and has a distinct physiological expression. Critically, we do not wish to claim that sleep states are identical to “local sleeps”, nor do we suggest an overlap between low arousal due to sleep deprivation and unconsciousness during sleep. To acquire estimates of their mean sleep schedule, participants wore an actimeter for one week before the total sleep deprivation protocol (Supplementary Fig. S1; available for 24/26 subjects due to data corruption). The total sleep deprivation protocol was as follows: A week prior to sleep deprivation, participants were provided with an actimetry device to track wake-sleep schedule, and were instructed to follow a consistent 8-hour sleep schedule. On the deprivation day, participants arrived at the lab one hour before their normal sleep time to extract their actimetry baseline data, estimate the optimal sleep deprivation window, and to provide baseline vigilance, drowsiness, and sleepiness measurements. After a total sleep deprivation of 26h (16h of typical wakefulness, 8h of sleep deprivation, and a 2h post-sleep deprivation period) participants began the post-sleep deprivation, experience-sampling session. As an example, a participant who typically slept at 12 am would arrive at the lab at 11 am, start sleep deprivation at 12 am, finish sleep deprivation at 8 am, and perform the experience-sampling task at 10 am. Should slow-wave activity during wakefulness follow the same circadian modulation it follows during sleep [50], a potential confound that could have lowered the power of our analysis is the time-window of the experience-sampling task. However, as suggested in [50], the relative time-window we have selected did not fall under a critical point of large reductions in the amplitude of the slow-waves. The 2-hour, post-deprivation waiting window allowed us to match the time of the experience-sampling across the 3 conditions, avoiding potential circadian confounds on experience-sampling, as we could easier match sleep-wake cycles and the time of the experience-sampling within each participant. We have chosen this sleep manipulation as similar manipulations have been previously used to examine the effects of sleep pressure [51, 52], and have been shown to elicit distinct low-arousal cortical profiles [53, 54], as well as changes in the sympathetic/parasympathetic balance [55].

Sleep deprivation was controlled with regard to light influence (illuminance = 15 lux during wakefulness and 0 lux during sleep), caloric intake (standardized meals every 4 h), and body posture (semi-recumbent position during scheduled wakefulness) to minimize potential masking effects on the sleep-wake regulatory system. Participants were not allowed to stand up except for regularly scheduled bathroom visits and did not have any indications of the time of the day. The experimenters continually monitored participants to keep them awake. In case of a sleep event, the experimenters first tried to awaken the participant through an intercom, and in case of failure, they manually awakened the participant. We also monitored for sleep lapses through the experience-sampling tasks. In case participants closed their eyes for a time period of < 30 seconds, they were probed by a tone to wake up. If they did not, the experimenter in the room would awaken the participant.

A one-week interval took place between sleep deprivation and further recordings in order to minimize potential carry-over effects of sleep deprivation on our follow-up conditions. In that way, the participants’ sleep schedules will also reset to their respective normal cycles. The order of the three arousal conditions was randomized. As a post-registration note, we randomized only the order of sleep deprivation and post-exercise, to add a training session before the baseline that allowed participants to get acquainted with the protocol, without external task impositions that might confound protocol understanding.

### Sampling Plan

We used a Neyman-Pearson frequentist approach to balance false-negative and false-positive rates by setting power to 95% and establishing a Type I error rate (alpha) of 5%. To estimate the desired sample size, a simulation approach was utilized: data were generated consistent with a latent binomial regression model, in which one categorical predictor with 3 levels (Base, High, Low) predicted a binary outcome Y (presence of MB or not). An original probability p_MB_ = 0.1 was specified as the underlying generative probability in the baseline model based on previous research [5, 11, 12]. We allowed the random intercepts and slopes to freely vary around a normal distribution with a standard deviation of s.d. = 0.1. Given that no previous study to our knowledge has provided evidence for the distribution of the effect sizes of arousal on mental reports, and to account for possible reverse effects (such as decreased MB report probability), we reasoned that a meaningful yet conservative effect for the *Low Arousal* condition would be an odds ratio of 1.6 and an odds ratio of 0.55 for the *High Arousal* condition. Since our initial hypothesized distribution is expected to yield ∼3-5 MB reports per session [11, 12], this effectively translates to a small effect size of interest of at least 3 more reports across conditions.

Considering these parameters, for each population sample, ranging from 5 to 50 participants, we sampled 500 datasets, and fit a binomial model with the participant ID as random factors, keeping the regression coefficients for the levels of the predictor constant. Based on the simulation analysis, using a false positive threshold of .05, we calculated a sample size of 26 participants to achieve a power of .95 (Supplementary Fig. S2).

## Data Analysis

### Behavioral data

Statistical analysis was performed using generalized linear mixed-effects models. To address whether arousal affects MB occurrence, we used a binomial, linear model with arousal as a categorical independent variable, and the proportion of mental reports across a sampling period (40 trials) as our dependent variable. Data were binary coded (presence or not of MB report) and fit into the model using a “logit” link. Given that the underlying distribution was unknown, a Bernoulli generative process minimized the assumptions about the model. In order to examine whether the multinomial distribution of mental reports itself changes across different arousal conditions, we used the generalized estimating equations (GEE) approach, an extension of generalized mixed-effects models that can account for correlated, repeated-measures count data from multinomial distributions [56, 57]. Mental reports were aggregated as counts across participants and conditions, and we examined shifts in report time distribution using the three experimental arousal conditions as predictors. We considered as report time the interval between the response probe and the participant’s report. To examine report times as a function of mental states, we specified a generalized linear mixed-effect model with mental reports and arousal conditions as categorical variables and used a gamma distribution with an “inverse” link function. As reaction times are usually an indicator of arousal effects on the task performance, an effect of arousal condition as a covariate might be informative about a potential shift of the overall slower mental report times distribution and about the arousal condition of the subject itself. The choice of the distribution and the link minimizes assumptions about the model, respects the positive, skewed distribution of reaction times, and was previously found to provide a better fit compared to other link functions [58]. To examine whether arousal shifts the dynamics of mental reports, i.e. one state might be more likely to be followed by MB in one of the arousal states compared to *Baseline*, we estimated dynamical transition probabilities across different mental states using Markov models. The transition probabilities for MB were then compared using a linear model with an identity link, with the transition probabilities as the dependent variable and the arousal condition as the categorical, independent variable.

All specified models were compared against null models using likelihood ratio tests. We introduced the participant’s ID as a priori random factor, i.e., we allowed the model’s intercept to vary. In case of multiple models compared, p-values were corrected using Bonferroni correction. In case of significance of a fixed predictor, we used corrected pairwise comparisons to examine the marginal means of the predictors.

### Brain-based measures

Physiological and cortical timeseries were segmented based on the response probe time. We considered the 110-second period before the response probe as a meaningful analysis epoch, representing the neuronal and physiological dynamics that result in a specific mental state. This period was used in subsequent analyses.

We recorded EEG with an EasyCap (64 active electrodes) connected to an actiCHamp system (Brain Products GmbH) using the 10-20 standard configuration. A ground electrode was placed frontally (Fpz in the 10–20 system). Online, we referenced the electrodes to a frontal electrode. Impedance was kept below 20 kΩ. As a post-registration note, we originally registered to keep impedance below 10 kΩ. However, we leveraged the strength of active electrodes and used the research standard of 20 kΩ. To minimize impedance, we used conductive gel. Data were sampled at a sampling frequency of 500 Hz. Preprocessing included band-pass filtering (0.1 Hz-45 Hz, FIR filter), notch filtering (50Hz), and epoch definition (t_start = 110s preceding the probe, t_max= probe). As a post-registration note, during EEG preprocessing, we observed low-frequency (<1 Hz) artifacts, such as sweat during the post-exercise session, that contaminated the quality of the signal. Therefore, we decided to reanalyze our data using a 1 Hz high-pass filter to minimize the presence of those artifacts. By visual inspection, we checked and removed noisy electrodes and epochs. In case of discarding more than 50% of the total epochs for a single participant, that participant was discarded from future analysis. We then used ICA decomposition to remove non-neuronal components such as blinks, heartbeats, muscle artifacts, etc. Finally, channels removed due to rejection were interpolated using neighboring channels, and all channels were re-referenced to the average.

Based on EEG recordings, we estimated three classes of measures: 1) measures estimating spectral power - raw and normalized power spectra, Median Spectral Frequency (MSF), spectral edge 90 (SEF90), and spectral edge 95 (SEF95), 2) measures estimating information content – spectral entropy, Kolmogorov-Chaitin complexity (K) and Permutation Entropy, and 3) measures estimating functional connectivity – Symbolic Mutual Information and weighted Symbolic Mutual Information. Power spectrum density (PSD) was computed over the delta (1-4 Hz), theta (4-8 Hz) alpha (8-12 Hz), beta (12-30 Hz), gamma (30-45 Hz) spectral bands, using the Welch spectrum approximation (segments = 512 ms, overlap = 400ms). Segment rejections were windowed using a Hanning window and zero-padded to 4096 samples. Kolmogorov-Chaitin complexity was computed by compressing a discretization of the signal using a histogram approach with 32 bins. Permutation Entropy was obtained by computing the entropy of a symbolic transformation of the signals, within the alpha, delta, and theta bands. SMI and wSMI were then computed from the same symbolic transformation, but data was first filtered using Current Source Density estimates to diminish the volume conduction. SMI and wSMI were computed in theta, delta, and alpha bands [59]. From the available connectivity metrics, we chose to use only wSMI as it is the only one that can detect purely nonlinear interaction dynamics and can be computed for each epoch [60].

### Physiological measures

<u>Electrocardiogram (ECG)</u> data were acquired using the BIOPAC MP160 system (BIOPAC SYSTEMS Inc.), amplified through the BIOPAC ECG100C amplifier. The data were sampled at a sampling frequency of 2kHz and recorded using the AcqKnowledge v4.4 software. ECG disposable adhesive skin electrodes were used in a bipolar arrangement of two electrodes and ground. The positive electrode was at the non-dominant wrist of the participant and the negative was on the contralateral ankle. The ground electrode was placed on the ipsilateral ankle.

ECG data were filtered with a notch filter (0.05 Hz) to remove baseline wander artifacts. A Butterworth high-pass filter was applied (0.5 Hz) to attenuate linear drifts and physiological artifacts. Powerline interference was attenuated with a notch filter (50 Hz). Finally, the data were smoothed with a 3^rd^-order polynomial Savitzky-Golay filter. Peaks were detected using the native Neurokit2 algorithm. Finally, data were epoched based on the partition scheme in the EEG preprocessing section.

ECG metrics were grouped into three domains: time, spectral power, and information content. Time-domain metrics were a) the Heart Rate (HR), b) the standard deviation of the RR intervals (SDNN), and c) the Root Mean Square of Successive Differences (RMSSD). Spectral power features were a) the Low Frequency of the Heart Rate Variability (LF-HRV), b) the High Frequency of the Heart Rate Variability (HF-HRV), and c) the LF/HF HRV ratio. Information content metrics were a) Approximate Entropy (AE), b) Sample Entropy (SE), and c) Multiscale Entropy (MSE). Initially, the native Neurokit2 algorithm to extract the peaks of the QRS complex. RR intervals were estimated as the sequential difference of the peak times. We estimated the time domain features based on the RR timeseries. For the spectral power metrics, the RR was evenly resampled at 4 Hz. Power spectra were computed over the LF-HRV (0.04–0.15 Hz) and the HF-HRV (0.15-0.4) bands. The power spectrums were estimated using the Welch procedure.

#### Respiration

Respiratory data was acquired using a respiratory belt and amplified through the BIOPAC amplifier. Data were sampled at a sampling frequency of 2 kHz and recorded using the AcqKnowledge v4.4 software.

Respiratory metrics were grouped in the time and information content domain. Time-domain metrics were a) respiration rate and b) respiration rate variability. Information content was estimated based on multiscale entropy.

#### Pupillometry

Eye movements and pupil size in both eyes were recorded using oculometric glasses (Phasya recording system) with a sampling frequency of 120 Hz. The eye tracker was calibrated at the start of each recording. Data was epoched based on the epoching scheme in the EEG preprocessing section. We identified 100ms blink periods around blinks and removed the whole segment, as pre- and post-blink periods can introduce pupil dilation artifacts while the eye is recovering to its standard size. We interpolated segments using 3^rd^-degree cubic interpolation. Dilation speed outliers were calculated by estimating the median absolute deviation (MAD) of each value. Samples exceeding the deviation threshold were removed. Pupil dilation was smoothed using a moving average filter and baseline-corrected with a 100ms period 2s after the probe.

Pupil metrics were grouped in the same three domains: time, spectral power, and information content. Time-domain metrics were: 1) Blink rate, 2) Pupil size, and 3) Pupil size variability. Spectral power metrics were: 1) Low-Frequency Pupil Component (LFC), 2) High-Frequency Pupil Component (HFC). The information content metric is MSE. The power spectra were estimated using the Welch procedure. As a post-registration note, we encountered issues extracting pupil metrics at the *Low Arousal* condition, as participants tended to have their eyes closed or partially closed for most of the trials. As our device was not sensitive to capture dilation in this setting, we additionally estimated a) Blink Rate, b) Blink Duration, c) Blink Rate Variability, d) Mean Eye Openness, e) Eye Openness Variability, f) Percentage of 70% Eye Closure and g) Percentage of 80% Eye Closure. As stated below, our registered plan was to reliably estimate all time, frequency, and complexity metrics that can be of use to our classifiers. Therefore, while we do not deviate from our original registered protocol, it is of note that these features could not be estimated reliably.

<u>Electrodermal activity (EDA)</u> data was acquired through skin electrodes on the index and middle finger and amplified through the BIOPAC amplifier. Data was sampled at a sampling frequency of 2k Hz and recorded using the AcqKnowledge v4.4 software. All EDA metrics originated from the time domain: a) Galvanic Skin Response (GSR), b) tonic EDA, and c) phasic EDA. Extraction of the phasic and tonic components of the EDA was conducted with deconvolution of the EDA signal with a biologically plausible impulse response function with initially fixed parameters that are iteratively optimized per participant [61].

### Pattern recognition

To examine the physiological counterpart of the behavioral shifts in MB reports, we employed a supervised decoding approach. Using the multimodal neurophysiological measurements during the three experience-sampling sessions, we trained multiple classifiers to discriminate across MB, MW, and SENS reports, to identify whether MB is supported by a unique brain-body interaction pattern. This approach allowed us to extract meaningful brain-body interactions from the proposed arousal metrics without being conservative about the nature of the multiple comparisons between the various body metrics.

As features, we opted to collect meaningful data in the time, frequency, information, and connectivity domain, unless such measurements could not be reliably estimated within our selected time window. The goal of the multiple selected metrics was to capture potential diverse spatiotemporal relationships (low-high frequency interactions, phase-amplitude interactions) that might extend across different recording modalities. Overall, we computed 57 features.

As targets, we used the participants’ mental states (MB, MW, and SENS). Since this creates a multiclass classification problem, we will focus on the binary classification of MB vs other reports. We expect to acquire 40 samples per participant and condition (i.e. baseline and arousal states), giving a total of 1040 (26*40) samples per condition. We expected that 5% of the samples correspond to the target report (MB), yielding an imbalanced problem with only 52 target samples per condition.

As learning algorithms, we tested parametric and non-parametric models, such as Support Vector Machines, Random Forests, and Extremely Randomized Trees. Support vector machines are a classification technique that aims to separate labeled inputs by creating a hyperplane that maximizes the distance of their features. Given a set of n-labeled inputs, SVM provides a hyperplane in an n-dimensional space that maximally separates the differently labeled groups. A random forest classifier is a meta-estimator. Various classifiers (“decision trees”) are trained in different parts of the input dataset, and each classifier uses only that part of the dataset to predict the label of the input. Then, the predictions of each classifier are pooled (“bagged”) together, and an optimal decision is chosen based on the label with the most predictions (“votes”). Finally, an extremely randomized tree classifier is a meta-estimator that employs a similar voting scheme. However, in the case of extremely randomized trees, trees are trained on all the features and the cutoff point of the trees (how the various metric nodes are arranged to reach a decision) is randomized. Since our problem is highly imbalanced, we also tested outlier detection algorithms (i.e. one-class classifiers), aiming to isolate MB from the other reports by considering MB as either an inlier or outlier. We then tested the one-class counterparts of the SVM (One-class SVM) and Random Forests (i.e. isolation forests) algorithms.

For model selection and performance estimation, we employed two different cross-validation approaches. First, we used a 5-fold stratified cross-validation scheme trained with all the samples. This provided us with performance estimates of classifiers aimed at obtaining patterns of brain and body function that can predict the report of MB in known participants. As a second approach, we used a 5-fold group stratified cross-validation scheme, using participants as groups. In this scenario, each participant was either on the train or on the test set. Thus, it aimed at learning general patterns of brain and body function that could predict the report of MB in unseen participants. In other terms, the first approach aimed at learning patterns that could discriminate MB from other reports while accounting for each participant’s variance, while the second strengthened the claim, aiming to learn general patterns that could be found in unseen participants.

As performance metrics, we report a) recall, b) precision, c) F1-score, d) area under the ROC curve (AUC), and e) balanced accuracy. Recall is the ratio of how often an item was classified correctly as a positive (True Positive / True Positive + False Negative). Similarly, precision is the ratio of actual correct positive classifications among positive classifications (True Positive / True Positive + Positive). F1-score is the harmonic mean of precision and recall. The AUC curve is another evaluation metric that summarizes how well the classifier predicts a class based on different thresholds of true positive and false positive ratios. Finally, balanced accuracy is an evaluation metric suitable for imbalanced datasets, where one class appears at significantly different frequencies than the others. Balanced accuracy is useful because it is estimated as the average of specificity and sensitivity, simultaneously controlling for very high precision due to classifying nothing as the infrequent class and very high recall due to classifying everything as the infrequent class.

We selected each model’s hyperparameters using nested cross-validation (same scheme as the outer cross-validation), using the F1-score as our optimization metric.

To evaluate the variance in the classifier performance and compare it to chance level, we performed repeated cross-validation (10 times), while training also a “dummy” classifier to obtain the empirical chance level of the training samples distribution. This type of classifier generates predictions based on the distribution of training samples for each class without accounting for the features.

The decoding analysis was implemented in Python using Julearn [62] and Scikit-Learn [63]. Metrics were estimated from existing Python libraries: MNE [64], NICE [65], Neurokit [66], and custom in-lab Python functions.

## Results

### Participants

To achieve a power of .95 at an alpha threshold of .05, we acquired 3 sessions of 40 trials per session from 26 participants (mean age = 26.38, std = 4.53, min=20, max=40; female=11). As a post-registration note, in case participants could not adhere to the strict 3-week protocol (30% total sessions), they were rescheduled to a later date that respected their sleep schedules to avoid time windows with potential extreme slow-wave activity [50]. Due to data corruption, one participant had 30 trials in one of the three sessions, and one participant had 33 trials in one of the three sessions. The remaining two sessions were completed for both participants.

### Behavioral Data

#### Occurrences of mental state reports alter across arousal conditions

We found a main effect for mental states, with MB being reported at significantly lower rates (Mean proportions ±SD: MW=.56, ±.21, SENS=.2±.14, MB=.12±.13; Kruskal H=124.07, p= 1.2e-27, eta^2^= .53) compared to MW (Dunn’s test=-10.75, p_FDR_ = 1.8e-26) and to SENS (Dunn’s test=-2.85, p_FDR_= 4.3e-03). Additionally, MW was reported significantly more frequently compared to SENS (Dunn’s test=7.9, p_FDR_= 4.3e-15; Fig. 2). As the study was focused on wakeful mental states, “SLEEP” reports were not included in the analysis (Mean proportions ±SD: *Baseline* = .03±.05, *High Arousal* = .05±.07, *Low Arousal* = .26±.21, Total = .1±.17).

**Figure 2.**
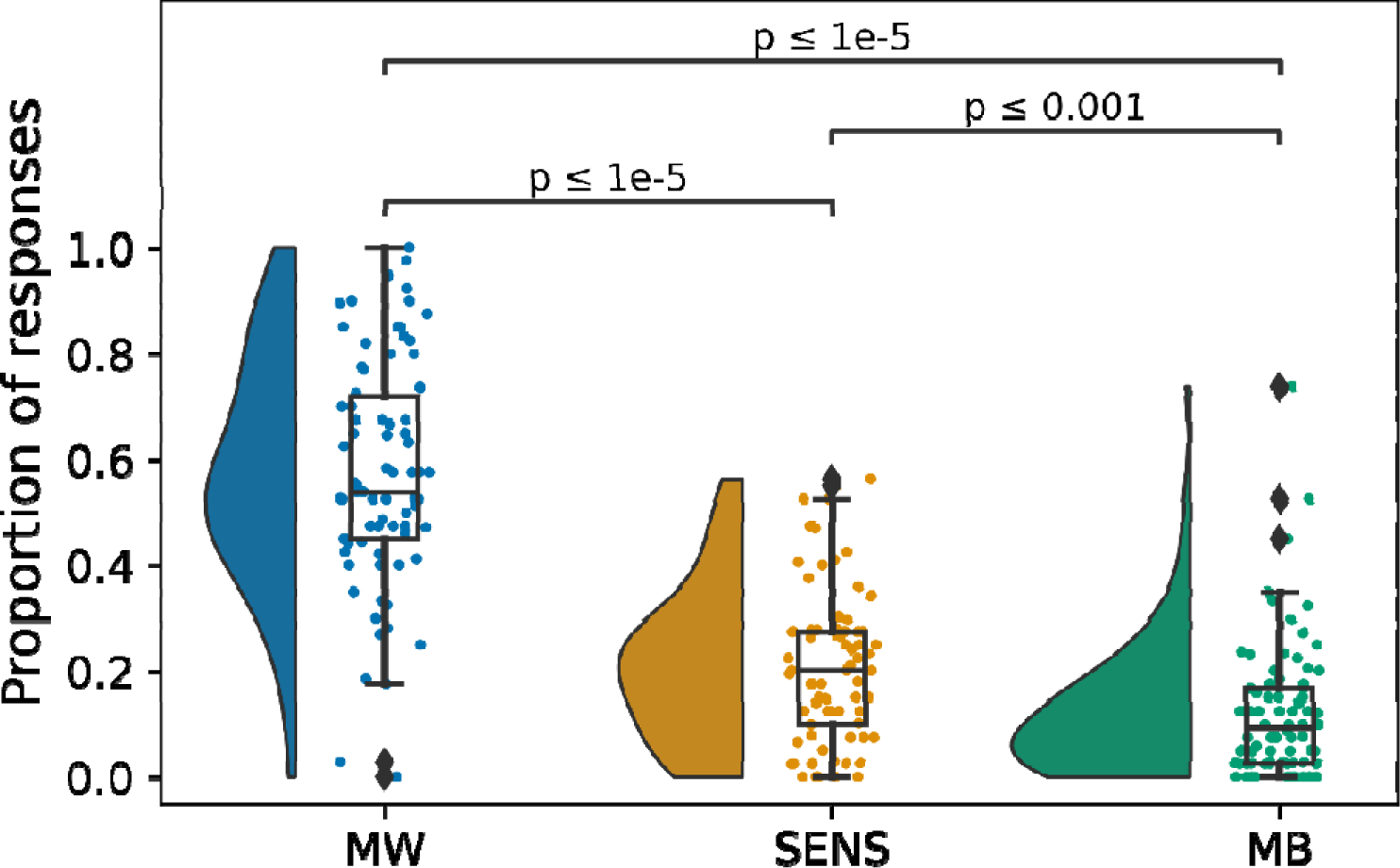
Mind-blanking (MB) was reported significantly less frequently compared to mind-wandering (MW) and Sensations (SENS) across all arousal conditions, validating what is generally reported in the literature. Density kernels show overall data dispersion and clustering trends. Point plots are individual subject estimates. Boxplots show medians and interquartile ranges, while whiskers indicate extreme values and diamonds indicate outliers.

We found that a model including all conditions outperformed a null model with only an intercept (Full_LogLik_ = -1021, Null_LogLik_ = -1046.83, χ^2^ = 51.57, df = 2, p_Bonf_ = 6.1e-12): MB was reported significantly more frequently in *Low Arousal* compared to *Baseline* (Marginal Mean= -.79, SE = .14, CL = [-1.16,-.43], p_FDR_ = 1.8e-08) and to *High Arousal* (Marginal Mean = -.97, SE =.15, CL = [-1.35,-.59], p_FDR_ = 7.9e-11) (Fig. 3a). However, MB reports during *Baseline* and *High Arousal* were comparable (Marginal Mean = .17, SE =.15, CL = [-.21,.56], p_FDR_ = 2.4e-01). A visual inspection of the individual marginal means showed that this effect was consistent across participants and was not driven by extreme cases (Fig. 3b-d).

**Figure 3.**
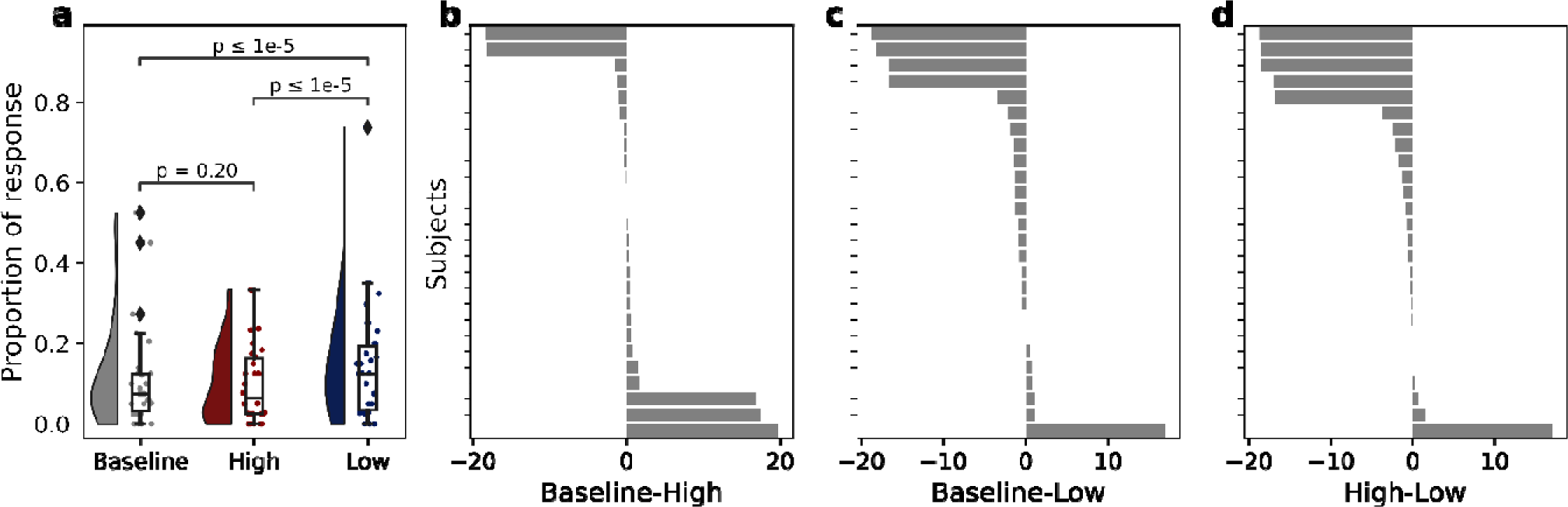
The frequency MB reports altered across the three arousal conditions. a) Mind-blanking (MB) report probability increased in *Low Arousal* (after sleep deprivation) compared to *High Arousal* (after intense exercise) and *Baseline*. Density kernels indicate overall data dispersion and clustering trends. Point plots represent participants’ MB report probabilities. Boxplots indicate medians and interquartile ranges, whiskers indicate extreme values, and diamonds indicate data outliers. b-d) Barplots denote single-subject marginal means, comparing MB reports across arousal conditions. Compared to *Baseline*, the e was no significant change during *High Arousal* (b). However, there was a visible trend favoring an increased probability of MB reports in the *Low Arousal* condition compared to baseline and *High Arousal*, signifying that the effect was present in most participants (c-d).

Additionally, generalized estimating equations (GEE) showed a significant interaction for MW between *Low Arousal* - *Baseline* (beta = 6, SE = 1.5, CL = [3.06, 8.94], p_FDR_ = 6.4e-05) and *Low* - *High Arousal* (beta = 8.23, SE =1.6, CL = [5.1, 11.36], p_FDR_ = 2.6e-07). We also found significant interactions in SENS reports, such that SENS tended to be higher in *Baseline* compared to *High* (*SENS Baseline - SENS High*: beta = 2.54, SE = .81, CL = [.96, 4.12], p_FDR_ = 1.7e-3) and *Low Arousal* (*SENS Baseline - SENS Low*: beta = 2.46, SE = .77, CL = [.96, 3.97], p_FDR_ = 1.3e-3). It is of note that this analysis yielded no significant results for MB, but the overall trend of the beta estimates was consistent with our positive results of the logit model above (Supplementary Fig. S3).

#### MB was characterized by higher reaction times

There was a main effect of arousal conditions, with reports during *Baseline* being reported the fastest and during *Low Arousal* the slowest (Fig. 4a). Also, there was a main effect of mental states, with MW reports being reported the fastest and MB reports the slowest (Fig. 4b). A significant interaction between MW and arousal showed that MW was reported the slowest in *Low Arousal* (Fig. 4c). A significant interaction between MB and arousal condition showed that MB was reported the slowest in *High Arousal* and *Low Arousal* (Fig. 4e). A model including both arousal and reaction times outperformed simplified models including only null or main effect terms (Full_LogLik=_ 2889.76, χ^2^ = 47.1, df = 4, p_Bonf_ = 1.5e-09; Fig. 4c). For a detailed overview of main effects and interactions, see Supplementary Table S3.

**Figure 4.**
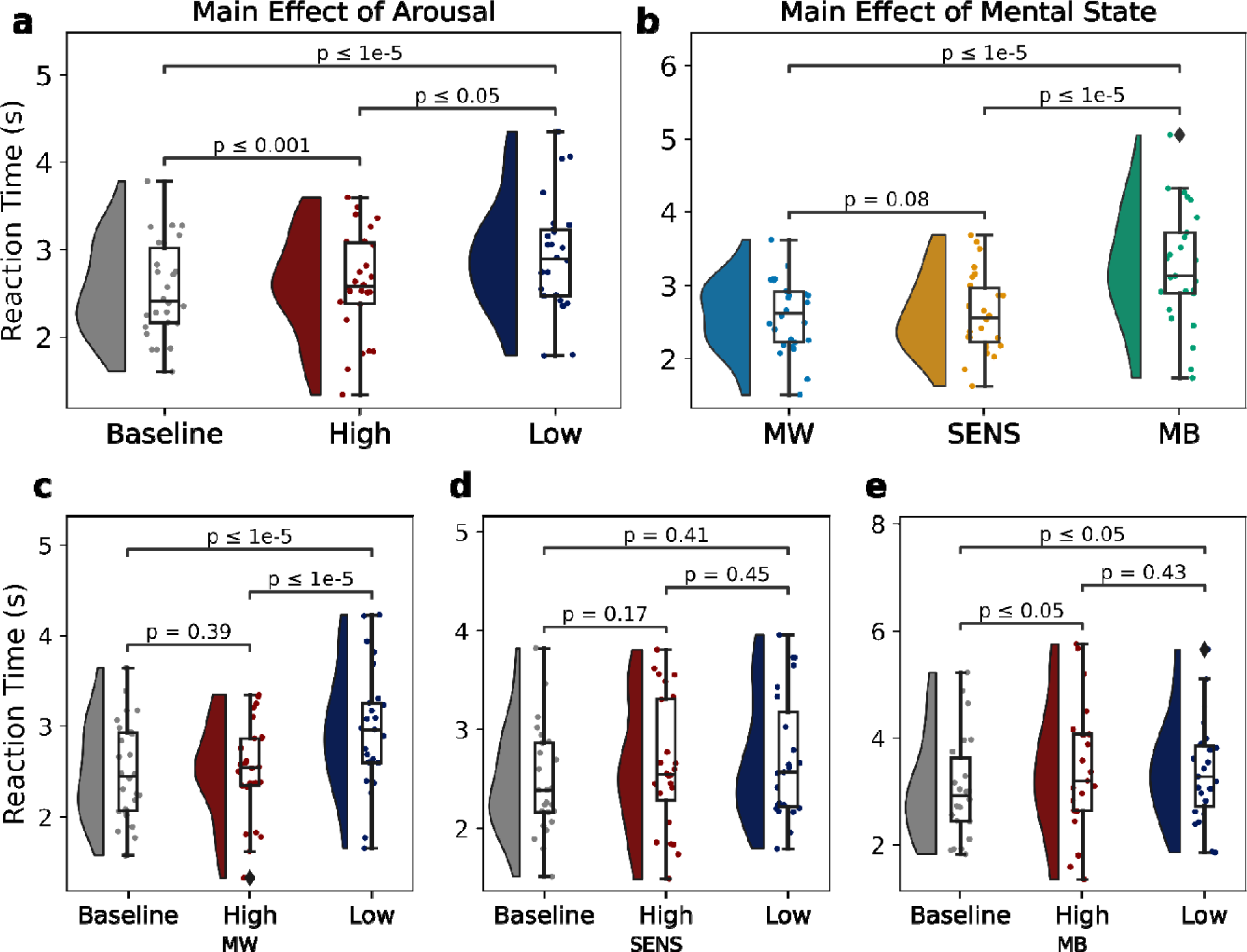
Mental states had different report times depending on arousal conditions. a) Reaction times at *Baseline* arousal were reported the fastest, followed by *High* (after exercise) and *Low Arousal* (after sleep deprivation), collapsed across all mental states. Pointplots show individual subject estimates. Boxplots show medians and interquartile ranges, while whiskers show extreme values. b) Mind-wandering (MW) was reported the fastest, followed by Sensations (SENS) and mind-blanking (MB), collapsed across all arousal conditions. Pointplots show individual subject estimates. Boxplots show medians and interquartile ranges, while whiskers show extreme values. c-e) Interaction between arousal condition and mental state report times: MW was reported the slowest in *Low Arousal* compared to *Baseline* and *High Arousal*, while MB was reported the slowest in the *Low Arousal* and *High Arousal* conditions compared to *Baseline*.

#### Transition probabilities showed reduced probability to transition to MW in *Low* arousal

Markov transition probabilities indicated significant differences only between *High* and *Low Arousal* conditions (Fig. 5), such that MW was more likely to be followed by MB (t = 3.26, CI = [.03,.15], p_FDR_= 9.7e-03, Cohen’s D = .74). Also in *Low Arousal*, both MW (t = -3.79, CI = [-.31, -.9], p_FDR_ = 7.6e-03, Cohen’s D = -.86) and SENS (t = -3.43, CI = [.37, -. 09], p_FDR_= 9.5e-03, Cohen’s D = - .77) were less likely to be followed by MW (Fig. 5; Supplementary Fig. S4).

**Figure 5.**
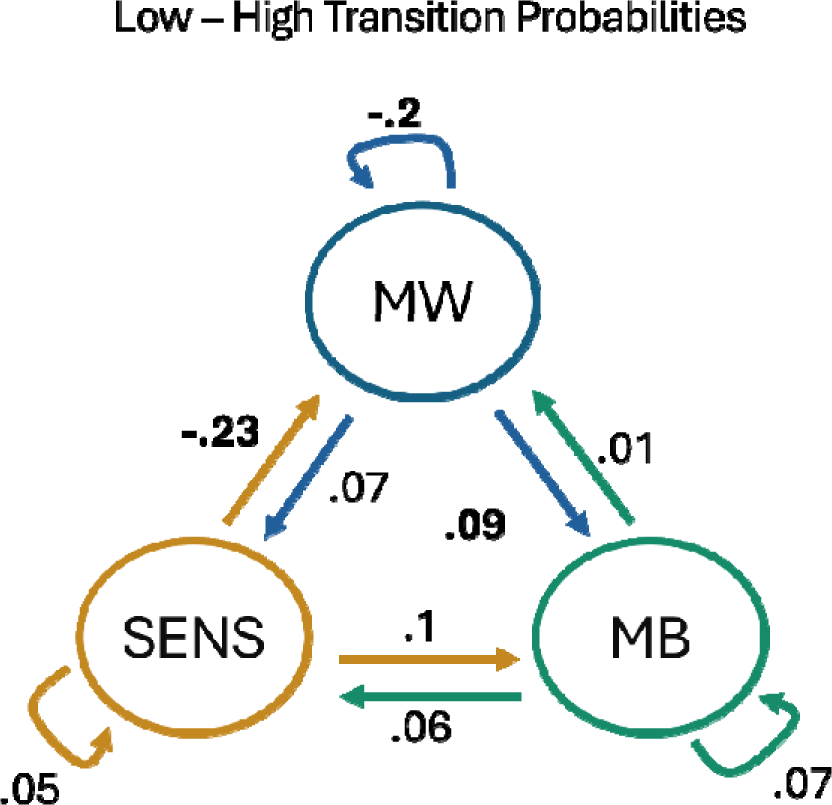
After sleep deprivation (*Low Arousal)*, participants were more likely to transition from mind-wandering (MW) to mind-blanking (MB) compared to the condition of physical exercise (*High Arousal)*. Additionally, participants were less likely to transition to MW. Arrows indicate the direction of the mental state transition. Bold font indicates statistical significance (FDR corrected).

#### Exploratory Analysis 1: MB frequency did not correlate with SLEEP frequency

As we wanted to avoid participants confounding MB and SLEEP reports, we opted for a paradigm that allowed participants to report both. Spearman correlations on each condition examined whether these two states were correlated. Across all mental states were comparable (*Baseline*: r = .13, p = 5.3e-01, *High Arousal*: r = .31, p = 1.3e-01, *Low Arousal*: r =-.05, p = 8.2e-01) (Supplementary Fig. S5). To strengthen the claim that MB and SLEEP reports do not covary, we additionally ran separate equivalence tests on each correlation. No test was able to reject an equivalence claim (*Baseline*: z = -.34, p = 3.7e-01, *High Arousal*: z = .54, p = 7e-01, *Low Arousal*: z = .72, p = 2.3e-01). Therefore, these results remain indeterminate.

#### *Exploratory Analysis 2: High Arousal* MB reports increased at the start, but not the end, of the experience-sampling session

While we found that MB reports were more frequent in *Low Arousal*, we did not find any significant effect of *High Arousal*. In our original hypothesis (Supplementary Table S1), we registered as a potential alternative explanation for the absence of an effect that high arousal, as elicited by high-intensity exercise, might not last for the full session, and our session would represent a gradual return to *Baseline Arousal*. To test for potential effects of more frequent MB reports only at the start of the experience-sampling we split the *High Arousal* session in two parts and compared the count of MB reports across the start and the end of the experiment. Using a chi-squared test we found a significant effect, with MB reports being more frequent (divergence = 4.08, p = 3.2e-02) during the first half of the *High Arousal* condition compared to the second half (MB_start_ = 93, MB_end_ = 66). We additionally attempted to validate this hypothesis by splitting the session into 4 and 6 discrete segments of 10 and 7 trials each and replicated the same analysis. However, this analysis did not reach significance. Finally, to provide further evidence for reduced occurrences of MB across time, we considered only the first and last 10 trials. We found a significant effect of more frequent MB occurrences (divergence = 7.39, p = 6.6e-03), with the first 10 trials of the *High Arousal* condition inducing more MB compared to the second half (MB_start_ = 51, MB_end_ = 27).

#### Classification of MB reports was outperformed by classification containing both BRAIN-BODY markers

We evaluated the capacity to classify MB reports from mental states with content (MW, SENS) based on 26 BRAIN (EEG) and 31 BODY features (12 ECG, 4 EDA, 8 RSP, 7 EYE), spanning time, frequency, information, and connectivity domains for each mental state report. In our original report, we registered that these features would be estimated across the 110s pre-probe window, with bad epochs being dropped. However, across an 110s epoch, even a nonlinearity of 1s would result in epoch removal, leaving a total clean sample of 25 / 78 sessions (29.4%), and a total of 1060/3120 (33.3%) clean epochs. Therefore, to preserve datapoints and data quality, and minimize data discarding due to brief non-linearities, we opted for an extra step in bad epoch removal. After the initial epoch definition of 110s, we followed it up by partitioning that epoch into 5s sub-epochs, resulting in 22 sub-epochs per epoch. We then proceeded to do bad epoch removal and EEG marker estimation on those sub-epochs. If an epoch consisted of more than 50% bad sub-epochs, it was discarded. Then, we averaged across within each epoch, resulting in no lost sessions, and a total of 2734 / 3120 (87.6%) total sample size.

Having a final 2734 reports x 57 features matrix per report, we trained multiple classifiers on the total dataset, to examine whether a specific brain-body profile would outperform chance level classification of MB reports (Table 1).

**Table 1.** A balanced random forest classifier outperformed all classifiers when compared across balanced accuracy. Cells indicate mean and 95% CI.

| Examined | Classifier | Recall | Precision | F1 | ROC AUC | Balanced Accuracy |
| --- | --- | --- | --- | --- | --- | --- |
| Known Subjects | Balanced RF | .62, [.6, .64] | .26, [.26, .27] | .37, [.36, .37] | .71, [.7, .72] | .66, [.65, .67] |
|  | SVM | .29, [.28, .31] | .28, [.27, .29] | .29, [.27, .3] | .62, [.61, .63] | .58, [.58, .59] |
|  | ET | .16, [.15, .17] | .61, [.58, .64] | .25, [.23, .26] | .73, [.72, .74] | .57, [.56, .58] |
|  | RF | .14, [.13, .15] | .57, [.53, .6] | .22, [.21, .23] | .71, [.7, .72] | .56, [.56, .56] |
|  | IF | .14, [.13, .16] | .2, [.19, .22] | .17, [.15, .18] | .52, [.52, .53] | .52, [.52, .53] |
|  | OC SVM | .89, [.86, .92] | .15, [.14, .15] | .25, [.25, .25] | .51, [.5, .51] | .51, [.5, .51] |
|  | DUMMY | .14, [.13, .15] | .14, [.13, .15] | .14, [.13, .15] | .5, [.49, .5] | .5, [.49, .5] |
| Unknown Subjects | Balanced RF | .46, [.41, .51] | .18, [.16, .2] | .25, [.23, .27] | .55, [.53, .57] | .54, [.53, .56] |
|  | IF | .23, [.19, .27] | .18, [.16, .2] | .19, [.17, .22] | .53, [.51, .54] | .53, [.51, .54] |
|  | RF | .05, [.04, .06] | .36, [.29, .44] | .08, [.06, .09] | .54, [.52, .55] | .51, [.51, .52] |
|  | OC SVM | .87, [.82, .92] | .14, [.13, .15] | .24, [.22, .26] | .51, [.5, .52] | .51, [.5, .52] |
|  | ET | .03, [.02, .03] | .36, [.26, .45] | .05, [.04, .06] | .53, [.52, .55] | .51, [.5, .51] |
|  | DUMMY | .15, [.14, .16] | .15, [.13, .17] | .14, [.13, .16] | .5, [.49, .51] | .5, [.5, .51] |
|  | SVM | .2, [.17, .22] | .16, [.14, .17] | .16, [.15, .17] | .49, [.47, .5] | .5, [.49, .51] |

RF = Random Forest; SVM = Support Vector Machine; ET = Extreme Trees; IF = Isolation Forest; OC SVM = One-Class Support Vector Machine

Due to the unbalanced nature of our dataset, we evaluated classifier performance based on balanced accuracy, as it avoids inflated performance on unbalanced datasets. Overall, we found that a balanced random forest (a random forest that undersamples the majority class in each bootstrap to equate class count) has above-chance performance and outperforms all other examined classifiers (Fig. 6a). We additionally examined whether we could predict unknown subjects, by leaving a subset of subjects out on each iteration. Due to the high degree of per-fold variance, we do not consider any classifier as meaningfully performing above chance level (Fig. 6b). Importantly, these results were replicated when we trained the classifiers in the 1-Hz filtered data (Supplementary Fig. S6a,b; Supplementary Table S4).

**Figure 6.**
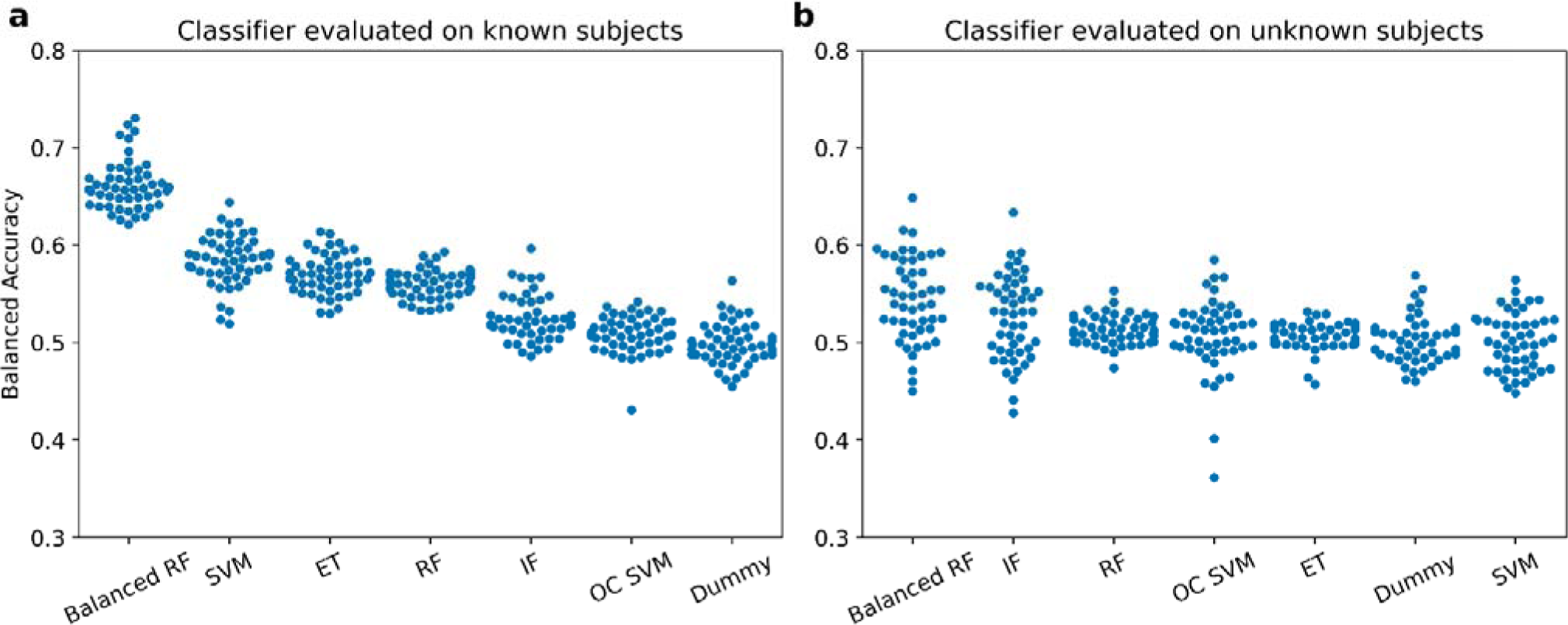
Classification performance was above chance level when mind-blanking (MB) reports were pooled across subjects, but not after training on a subset of participants and classifying the remaining subset. a) A balanced random forest classifier provided the highest classification performance across all examined classifiers including known subjects. b) An isolation forest classifier provided the highest classification performance across all examined classifiers on unknown samples. However, due to the high variance, we could not consider it meaningful. Individual points indicate performance on the folds of the repeated cross-validation. Results are ordered based on descending order of performance. Chance level performance is indicated by the Dummy classifier. RF = random forest; SVM = support vector machine; ET = extreme trees; IF = isolation forest; OC SVM = one-class support vector machine.

Having established that MB reports can be predicted from known subjects, we then examined whether a brain-body data pattern would outperform classifiers trained solely on either BRAIN or BODY features. To this end, we fit and optimized a separate balanced random forest classifier on discrete feature combinations of our dataset. For a full report of the performance on different features, see Table 2 and Supplementary Table S5.

**Table 2.** A classifier trained on a combination of BRAIN and BODY features outperformed classifiers trained solely on BRAIN or BODY features, when evaluated with balanced accuracy. Cells indicate mean and 95% CI.

| Classifier | Recall | Precision | F1 | ROC AUC | Balanced Accuracy |
| --- | --- | --- | --- | --- | --- |
| BRAIN + BODY | .62, [.6, .64] | .26, [.26, .27] | .37, [.36, .37] | .71, [.7, .72] | .66, [.65, .67] |
| BRAIN | .61, [.59, .62] | .24, [.24, .25] | .35, [.34, .36] | .7, [.69, .71] | .65, [.64, .65] |
| BODY | .59, [.58, .6] | .22, [.21, .22] | .32, [.31, .32] | .66, [.66, .67] | .61, [.61, .62] |
| EYE | .57, [.55, .59] | .21, [.21, .22] | .31, [.3, .32] | .64, [.63, .65] | .61, [.6, .62] |
| ECG | .55, [.54, .57] | .18, [.17, .18] | .27, [.26, .27] | .58, [.57, .59] | .56, [.55, .57] |
| EDA | .6, [.57, .63] | .17, [.17, .17] | .26, [.26, .27] | .57, [.56, .58] | .55, [.54, .56] |
| RSP | .52, [.5, .54] | .15, [.15, .16] | .24, [.23, .24] | .53, [.52, .54] | .52, [.51, .53] |

Overall, we found that a classifier trained on both BRAIN and BODY markers marginally outperformed classifiers trained solely on BRAIN or BODY features across all our performance metrics (Fig. 7a,c; Supplementary Fig. S7a,c; Table 2; Supplementary Table S5). To evaluate the impact of the number of features on the capacity of the learning algorithm to extract relevant information, we also trained the balanced random forest model using randomly shuffled bodily features. EEG features were not altered. The model with the shuffled values showed a decline in classification performance, providing evidence that, when classifying mental states, a model trained on both brain and body data learns unique information from both domains (Fig. 7d; Supplementary Fig. 7d). For feature importance, we calculated SHAP values for each feature in our dataset. SHAP values estimate the marginal contribution of each feature, averaged across every potential feature combination. In this manner, each value represents how much this feature contributes to the classification, after controlling for the impact of other features on this feature’s importance. We found that the model relied mostly on EEG and EYE openness features to discriminate MB reports when pooling MB occurrences across all three conditions. (Fig. 7b; For an extensive list of all SHAP values, see Supplementary Fig. S8). Importantly, feature importance did not substantially change when filtering the data with a 1 Hz filter (Supplementary Fig. S7b; For an extensive list of all SHAP values, see Supplementary Fig. S9). Overall, the comparable performance of the models, and the high degree of overlap in the ranking of the feature importance point to the robustness of the models.

**Figure 7.**
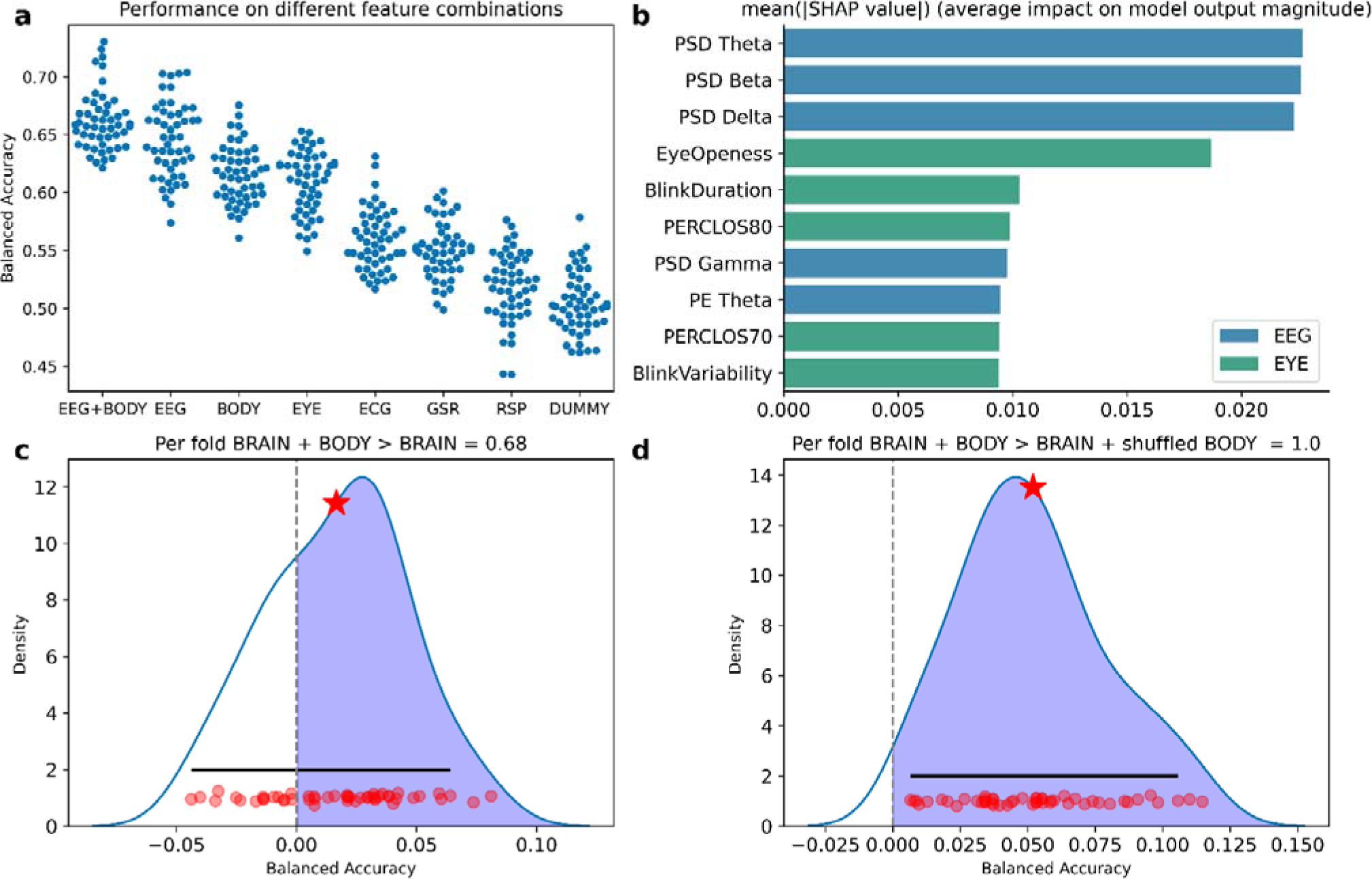
Classification of MB improves when considering both BRAIN and BODY. A balanced random forest classifier trained on a combination of BRAIN and BODY features outperformed classifiers trained solely on BRAIN or BODY features when evaluated with balanced accuracy. Individual points indicate performance on the folds of the repeated cross-validation. b) Subset of the 10 features with the highest mean of the absolute SHAP values obtained from the balanced random forest classifier. c) The per-fold differences between the classifier trained on both BRAIN and BODY features and the one trained only on BRAIN data suggest that incorporating both feature domains provides a slight performance improvement over using BRAIN data alone. The shaded region indicates better performance for the classifier trained on both feature domains. The star indicates the mean difference. The solid, horizontal line represents the 95% highest-density intervals of the distribution. Red dots indicate per-fold differences. d) The per-fold differences between the classifier trained on both BRAIN and BODY features and the one trained on BRAIN and shuffled BODY data suggest that the model with both BRAIN and BODY data does not consider the body markers as noise.

#### Exploratory analysis 3: Feature importance altered across arousal conditions

The decoding analysis in known samples showed that we can predict MB instances from the combination of brain-body markers with adequate accuracy when MB instances were aggregated across different arousal conditions. We were further interested in whether this classification was achieved based on a universal mechanism, or whether we could detect arousal-dependent brain-body configurations that predict MB. To this end, we trained a balanced random forest classifier solely on data acquired from *Baseline*, from *High,* and from *Low Arousal*. We found that *Baseline* had the best performance (.67, [.65, .68]), followed by *Low Arousal* (.64, [.63, .65]), and finally *High Arousal* (.61, [.6, .63]). We retained comparable performance when examining the arousal partitions of the 1 Hz filtered dataset (Supplementary Table S6-7). Examining the SHAP values for each arousal state, we saw that the models relied on distinct feature domains. During *Baseline*, the model relied on markers from the frequency domain of EEG (Fig. 8a). During *Low Arousal*, MB classification was obtained using the delta band power, by far the most dominant marker (Fig. 8b). Finally, in *High Arousal*, the model did not rely on a single feature, rather in a combination of eye openness, GSR, and the frequency domain of EEG (Fig. 8c). Similar feature importances were observed in the 1Hz filtered dataset (Supplementary Fig. S10). However, in the 1 Hz filtered dataset, we observed that ECG features tended to rank higher (Supplementary Fig. S11-16).

**Figure 8.**
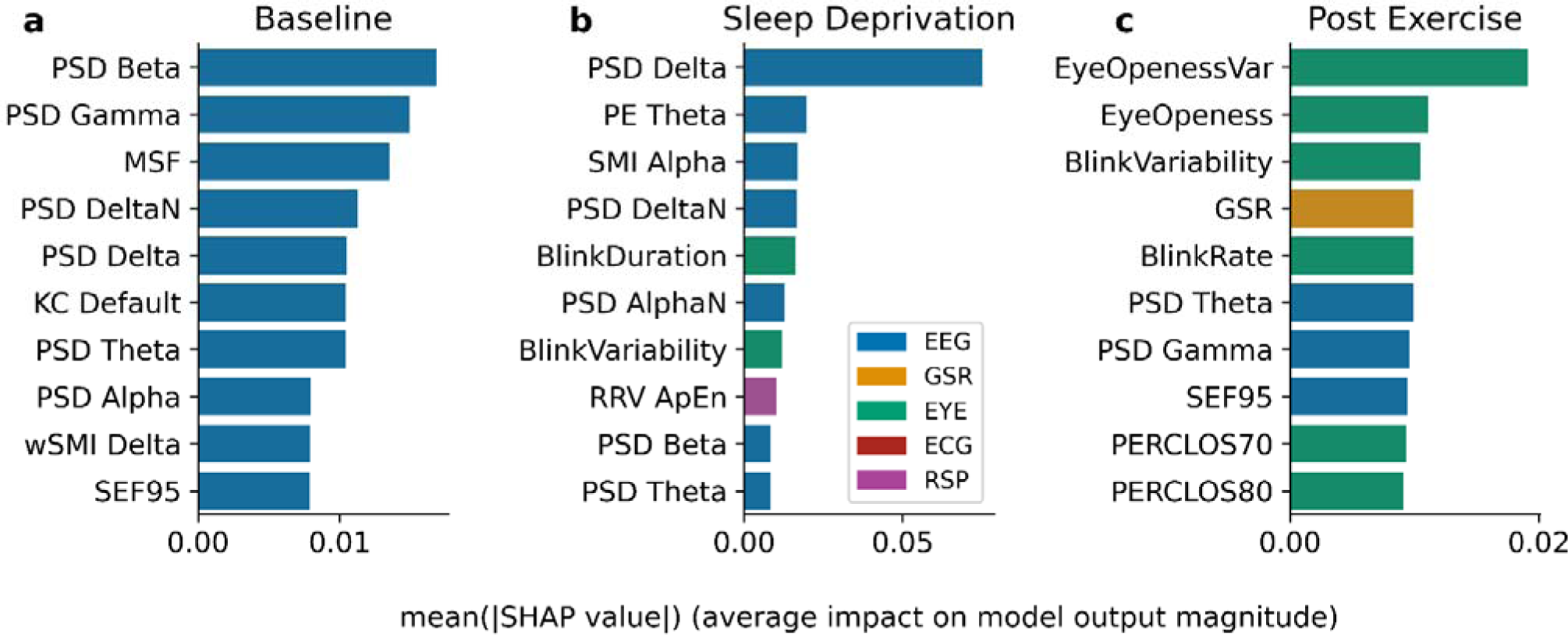
Ranking of features by mean absolute SHAP value extracted from the balanced random forest classifier varied across different arousal conditions. a) Magnitude of SHAP values for a balanced random forest classifier trained on MB reports collected during the *Baseline Arousal condition*. The model relied mostly on features from the EEG frequency domain. b) Magnitude for SHAP values for a classifier trained on MB reports collected during the *Low Arousal* condition (after sleep deprivation). The model mostly used spectral power in the EEG delta band. c) Magnitude for SHAP values for a classifier trained on MB reports collected during the *High Arousal condition* (after intense exercise). The model relied mostly on features from eye openness, EDA, and the EEG frequency domain.

#### Exploratory analysis 4: Feature importance altered based on the pre-probe analysis window

A potential caveat of utilizing the full pre-probe period of 110s before a report is that we might capture multiple mental states, and the actual statistical regularities might be weakened when averaged across. With this consideration, we examined whether we could improve classification performance when classifying MB from the last 10s before a report. We defined a secondary brain-body data matrix, with body features that could be estimated from 10s of body activity. Across both 0.1 and 1 Hz filters we retained comparable performance in the classifiers trained on both EEG and bodily markers, as well as solely EEG or body markers (Supplementary Fig. S17-20; Supplementary Table S8-9). However, we observed decreased performance in the classifier trained solely in the eye openness data (Supplementary Table S8-9). An examination of feature importance showed that the beta, delta, and theta bands of the EEG frequency domains remained the most important EEG features, but there was a reduction in the importance of the EYE features and an increase in the importance of EDA (Supplementary Fig. S17b,18,19b,20). Importantly, our results were not affected by the choice of filtering parameters, indicating robustness of our results to preprocessing parameters.

## Discussion

We used experience-sampling combined with EEG and peripheral physiological recordings under different autonomic arousal conditions to determine whether MB reports in neurotypical individuals were supported by distinct brain-body configurations compared to mental states with reportable content. Overall, our results show that MB is a mental state that becomes more prevalent in low and partially in high arousal states, and that MB is driven by both brain and body processes, providing evidence for an embodied account of MB.

Behaviorally, we found that MB was reported at significantly lower rates compared to sensory experiences or MW, irrespective of the arousal condition. This finding is in line with past research showing that MB rates vary between 5-10% of total probe instances, across both uninterrupted thinking [12] and task engagement [11]. We also show that sleep deprivation significantly increases the frequency of MB occurrences. Sleep deprivation induces a low arousal state during which cognitive performance declines [67], metabolic and physiological processes change [68], and unique neuronal markers, such as slow-wave activity, emerge [69]. After sleep deprivation, participants also tend to perform worse in sustained attention tasks [70], with results suggesting a true effect of sleep deprivation on more “misses” (no response when necessary) compared to “false alarms” (response when unnecessary) [71], a finding that was recently shown as a behavioral correlate of MB [11]. Additionally, sleep deprivation and mounting sleep pressure were positively correlated with more MW instances [72, 73], suggesting an overall mode shift from task engagement to MW [74]. Our results challenge these past findings by showing that participants were more likely to blank than mind wander after sleep deprivation. We also show that MW was in fact more likely to decrease after sleep deprivation. This is further supported by the results of the transition matrix analysis, where MW reports were less likely to be followed by another MW report, and more likely to be followed by MB. Such discrepancies in the reportability of MW after sleep deprivation could be possibly explained by the explicit inclusion of MB as a reportable mental state in the experience-sampling that our design opted for. In other words, it might be that the observed MW occurrence increase after sleep deprivation could be accounted for by MB reports, once participants had the chance to opt between these two mental states in a more fine-grained way.

In terms of high arousal induced by high-intensity exercise, our analysis did not reveal any significant effects on MB occurrences. As per the provided registered protocol alternative explanation (Supplementary Table 1), we hypothesized that this arousal manipulation might not have been overall effective as it could not produce effects that would last across the whole experience-sampling session. To test whether MB frequency reports would differ between the beginning and at the end of the session, we split the dataset into two parts. When split, we indeed found a significant difference between the frequency of MB reports. This result was replicated when considering only the first and last 10 trials per subject, which maximized the distance between initial and final physiological arousal within the session. However, we were not able to find any differences when the data were split into smaller bins. Together, we consider that these results provide partial evidence for our registered hypothesis, showing that residual high arousal effects after intense exercise can increase the frequency of MB reports.

In addition to the frequency of mental states across arousal conditions, we also examined whether report times differ across arousal conditions and mental states. In general, reports in low arousal tended to be the slowest, consistent with a wide range of attention tasks that show slower report times in sleep deprivation compared to baseline arousal [75]. We consider these findings as additional evidence that the arousal manipulation was effective in that it lowered overall vigilance levels. We also observed a main effect of mental states, such that MB tended to be reported significantly slower compared to MW and SENS. Contrary to our current results, we recently found that MB was reported faster when compared to other mental states when content had to be evaluated [12]. This apparent mismatch in results can be explained when considering that MB can be a state devoid of content, and therefore, there is the binary consideration of “yes/no” when evaluating thought content, which might be a relatively fast decision. This can be different, for example, from the evaluation of content-full mental states, which demand a sequential evaluation of both content presence (“yes/no”) and content evaluation (“what is the content about?”). This way, the difference in results can be explained by the imposition of an additional cognitive evaluation. Overall, we suggest that these results might reflect a gradient of vigilance, with participants being the most alert at baseline arousal, and progressively declining during high and low arousal conditions, as well as more vigilant when reporting mental states with content compared to MB. Of note, we observed two interesting interactions between mental states and arousal conditions. MW tended to be reported slower in low arousal compared to baseline and high, which is consistent of our interpretation of reaction times as marking vigilant states. However, as we also observed that MB reports tended to be reported the slowest in both High and Low arousal conditions, we speculate that this might be preliminary evidence that arousal modulates how engaged participants are with their current mental states. In this sense, exercise fatigue can lead to an MB state that takes longer to recover from when probed for a report.

A final explanatory analysis revolved around the relationship between sleep and MB. We recently posited that MB is a distinct mental state characterized by a unique phenomenological profile of no content [76], and unique neuronal markers, characterized by high cortical integration and low cortical segregation [12]. This neuronal configuration is atypical of wakefulness [77], and is more closely reminiscent of brain configurations during deep sleep [78]. In conjunction with the presence of slow wave intrusions during wakefulness as a marker of MB reports [11], a classic marker of NREM sleep, an emerging issue is whether MB is a misrepresented instance of sleep. This issue is further complicated by the postulation that in MB there is no content [76], and thus does not functionally represent a wakeful state where people can recover content. To avoid this pitfall, we introduced *Sleep* as a potential response during experience-sampling. We found that people discretely reported MB and sleep, providing evidence that when provided with such options, people can differentiate between these two experiences. Additionally, we did not find that MB and sleep tended to covary. To strengthen this claim, we ran equivalence tests for each correlation across arousal conditions. However, no test showed a positive result for equivalence. Therefore, these results remain indeterminate, with a trend for no relationship between MB and sleep.

Having established that MB occurrence varied across different physiological arousal conditions, we then examined whether MB could be decoded by brain and body markers. With the aim of showing single trial prediction, we trained different models on EEG and physiological signal markers from time, spectral, complexity, and connectivity domains. Overall, we were able to achieve above-chance-level classification, showing that there exist unique brain-body patterns that can discriminate MB reports from mental states with content. However, we were not able to show above-chance-level classification when training classifiers on unknown subjects. Therefore, our results are not generalizable to novel populations due to the high amount of variance between subjects. Of importance is the result that a combination of EEG and physiological markers marginally, but consistently outperformed both EEG and physiological markers. Overall, we observed an improvement of 2-5 % in classification performance in balanced accuracy. This improvement can be attributed to unique information inherent in body signals, as evidenced by the comparison of the classifier trained on both brain and body data compared to classifiers trained solely on brain data or brain and shuffled body data. The classifier trained on both brain and body data does not consider body features as noise or redundant. Overall, while our results suggest a high degree of overlap between brain and body information in MB, they indicate that information about MB extracted from the body is partially independent of the EEG features. Feature importance ranking derived from the decoding model indicates that the low and mid frequencies of the EEG power spectrum and metrics of eye openness are useful predictors of MB. This finding was consistent across analysis windows and preprocessing parameters. Importantly, all classifiers trained on body markers had above chance performance with variant degrees of variability, with the highest performing being the EYE (eye openness) and the ECG (heart-rate variability), providing evidence that MB can be decoded solely from bodily signals.

To further validate our protocol, we ran two exploratory analyses, with the aim to examine whether classification performance varies based on the analyzed pre-probe window and whether feature importance alters across arousal conditions (For a full Discussion, see Supplementary Discussion on Methodology). Overall, when examining a classifier trained on a brief 10s window before MB reports, we found comparable performance compared to the full 110s classifier. What was interesting was that while EEG performance remained the same, performance on classifiers trained solely on body features decreased. As brain-physiology coupling occurs at varying time delays across cardiac [79] and respiratory domains [80], we interpret these results as evidence that bodily contributions on MB are based on slow, oscillatory processes that might not be captured from examining short pre-probe periods. At the same time, our classification analysis on separate arousal conditions showed distinct brain-body configurations that can predict MB reports. As our decoding approach does not permit any inference of the directionality effect, or decomposing interactions within and across physiology modalities, at this stage we claim that our results point to discrete physiological pathways that elicit MB reports. Overall, we show that our enhanced classification is retained across different analysis windows and different arousal conditions.

Similarly, enhanced classification when considering a brain-heart matrix compared to solely brain markers was also shown for patients with disorders of consciousness, where the inclusion of cardiac features outperformed classification based solely on EEG markers [81]. To our knowledge, our results are the first to extend multivariate decoding past the brain-heart axis and consider the inclusion of multiple unique bodily afferent sources in classifying mental states. The overall success of the brain-body decoding paradigm in classifying consciousness levels and mental states provides evidence that bodily information is not redundant and is not necessarily fully represented within brain dynamics. Instead, an embodied approach, stressing bidirectional information routes between brain and body can provide better predictive power and assist in more comprehensive generative, computational models of experience [34, 82].

A neurobiological explanation of our results comes from an integrative model about content, task engagement, and arousal which suggests that the relationship between thought and arousal can be conceptualized as an inverted u-curve. This means that an optimal arousal level modeled by LC-NE firings is necessary to actively engage and control our thoughts, either during task engagement or MW [83]. This stance treats thought as an active task, where engagement is necessary for clear content and control of thought dynamics. As arousal tapers off to non-optimal levels of the inverted u-curve, we experience concurrent, opposing thoughts that serve exploratory purposes for optimal performance, such as exploring different strategies. This necessitates flexibility and malleability of content. We here suggest that our results supplement this model by providing an account of the extremities of the optimal u-curve. As the model suggests degradation of thought clarity when we move closer to arousal extremities, we consider MB reports as instances where no content can be clear or present, extending this unifying framework to the entire arousal u-curve. Neurophysiologically, this model has translated to investigations of pupil dilation, an index of LC-NE firing, as a function of mental state and task engagement with pupil size yielding both positive [26, 84] and null results [11] in discriminating on-task vs off-task mental states, as well as contrasting MB and MW. Part of the ascending arousal network, the LC modulates cardiac, galvanic, respiratory, and pupillary activity [28, 85]. In addition, the LC innervates projections responsible for eyelid openness [86]. The combinatorial high performance of different body markers in classifying MB reports, and the evidence that altered levels of arousal increase MB occurrences provide further support for the modulatory role of the ascending arousal system in mental states and thought reportability.

From a theoretic perspective, our study challenges the conception that brain information is uniquely suitable to understand thought reportability and provides support for an embodied account of the mind. Embodiment moves the seat of mental events away from the brain and reformulates cognition as resulting from brain-body interactions. An extensive literature has shown how cataloged cardiac, respiratory, gut, and pupillary effects on perception [30], action [87], metacognition [31] and consciousness [81], while the collective interplay of peripheral systems has discriminatory power for clinical [88] and consciousness classification [89]. We show here that within embodiment, the body is not only facilitatory but also might impede access to our mental lives. Under specific brain-body configurations, we are not able to clearly formulate mental content.

Some limitations pertain to our study. First, the nature of experience-sampling discretizes the continuous nature of ongoing thinking. As there is no consensus as to how long a mental state might last, or whether all mental states last the same length, results might average across different mental states. While we attempted to circumvent this problem by analyzing different pre-probe windows, it remains unclear whether all mental states last the same, and what is their actual duration. Secondly, the post-exercise setup might be suboptimal in examining the effects of high arousal on ongoing cognition. Neuronal and electrophysiological recordings have shown that the duration of the effects of exercise on ongoing brain and physiological activity [45, 46, 47] is highly variant. In addition, it is unclear whether brain and body recover to baseline states at the same rates, potentially confounding the post-exercise importance of cortical and physiological markers in cognition. Experience-sampling with online probes during exercise could overcome this challenge.

In conclusion, our study suggests that MB is an arousal-modulated mental state, with a unique cortical and physiological profile. We think that our results pave a new paradigm for an embodied account of mental states, where the phenomenology of our mental lives is expressed based on both our body and our brain state. At the same time, our results challenge the neurocentric approach to mental state research, putting emphasis on the constant brain-body interactions that shape our cognition. As MB research continues to evolve, we consider our findings elaborative for clinical and experimental accounts of the mind, where we move towards a complex and dynamic conception of our mind.

## Code Availability

All codes to replicate the power analysis, the experience-sampling paradigm, and the present analysis can be found at https://gitlab.uliege.be/Paradeisios.Boulakis/mind_blanking_arousal. An archived version of the code at the time of submission can be found at https://doi.org/10.58119/ULG/174Q6G.

## Data Availability

The aggregated raw data in a BIDS format, the trained machine-learning models, experimental and analysis logs, and result dataframes can be found at https://doi.org/10.58119/ULG/174Q6G.

## Protocol Registration

The stage 1 accepted-in-principle protocol can be found at https://osf.io/sh2ye. The authors confirm that no data for the pre-registered study was collected prior to the date of AIP.

## Acknowledgments

The experimental work was conducted at the GIGA-CRC-Human Imaging platform of ULiège, Belgium.

## Funding

At the time of the research, PAB and SM were FNRS Research Fellows. AD and CS are FNRS Research Associates. This work was supported by the Belgian Fund for Scientific Research (FRS-FNRS), the European Union’s Horizon 2020 Research and Innovation Marie Skłodowska-Curie RISE programme NeuronsXnets (grant agreement 101007926), the European Cooperation in Science and Technology COST Action (CA18106), the Léon Fredericq Foundation, and the University and of University Hospital of Liège. The funders had no role in study design, data collection and analysis, decision to publish, or preparation of the manuscript.

## Author Contributions

**Paradeisios Alexandros Boulakis**: conceptualization; data curation; formal analysis; investigation; methodology; project administration; software; visualization; writing – original draft preparation. **Nicholas John Simos**: investigation; software; validation; writing – review and editing. **Zoi Stefania**: investigation; project administration; writing – review and editing. **Sepehr Mortaheb**: formal analysis; software; writing – review and editing. **Christina Schmidt**: methodology; writing – review and editing. **Federico Raimondo**: formal analysis; methodology; software; validation; supervision; writing – review and editing. **Athena Demertzi**: conceptualization; methodology; supervision; writing – original draft preparation.

## Competing Interests

The authors declare no competing interests.

## Notes

### Competing Interest Statement

The authors have declared no competing interest.

### Summary of Updates

The revised manuscript contains supplementary analyses that ensure the machine-learning models' robustness across different preprocessing parameters.

