## Supplementary Material for "Variations of autonomic arousal mediate the reportability of mind-blanking occurrences"

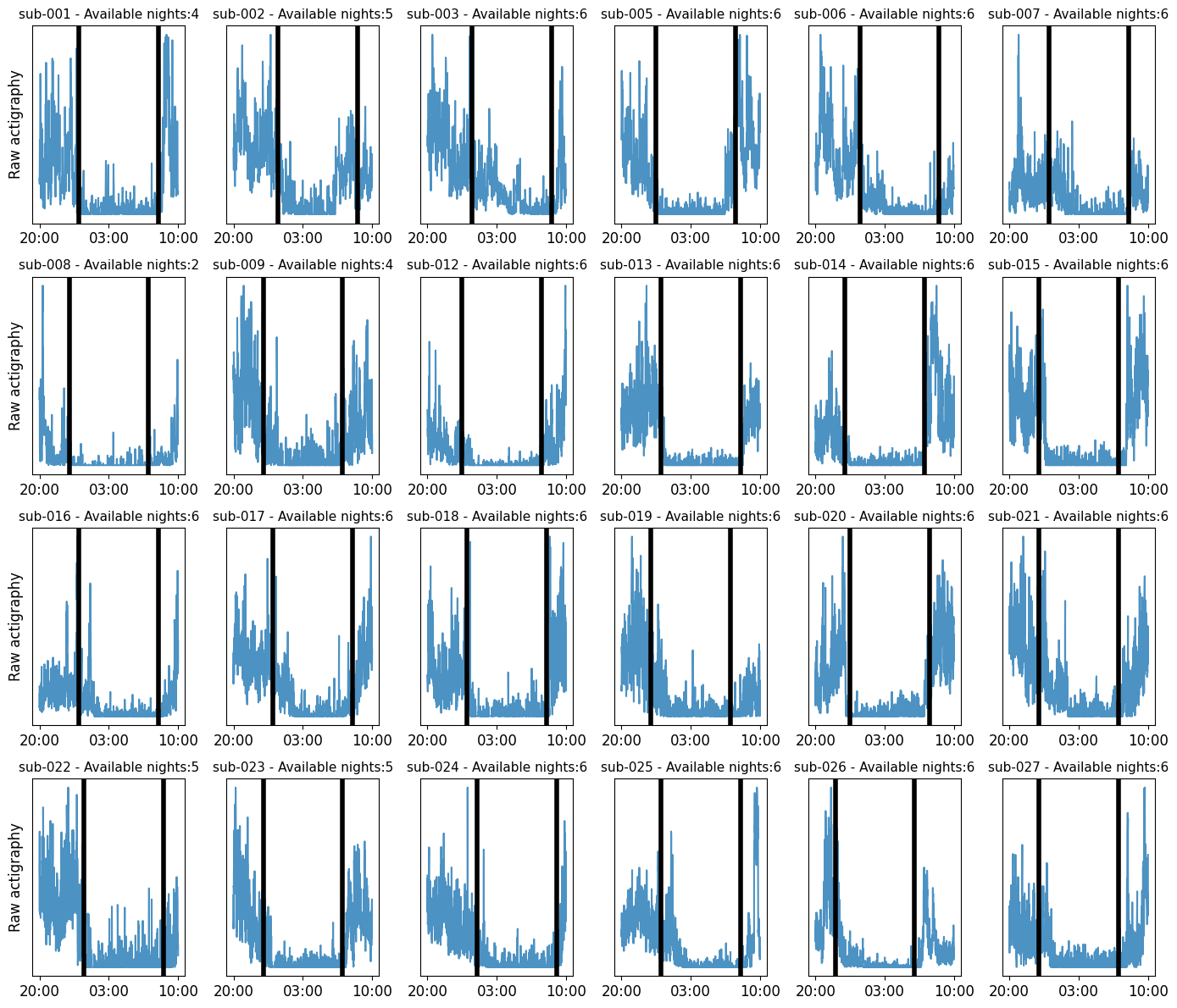

**Supplementary Figure S1.** Raw actigraphy plots (available for 24/26 subjects). We observed a reduction in actigraphy activity in the allocated sleep windows, indicating that participants maintained a steady sleep schedule in the week preceding the sleep-deprivation session. Vertical black lines indicate sleep onset and sleep end.

**a)**

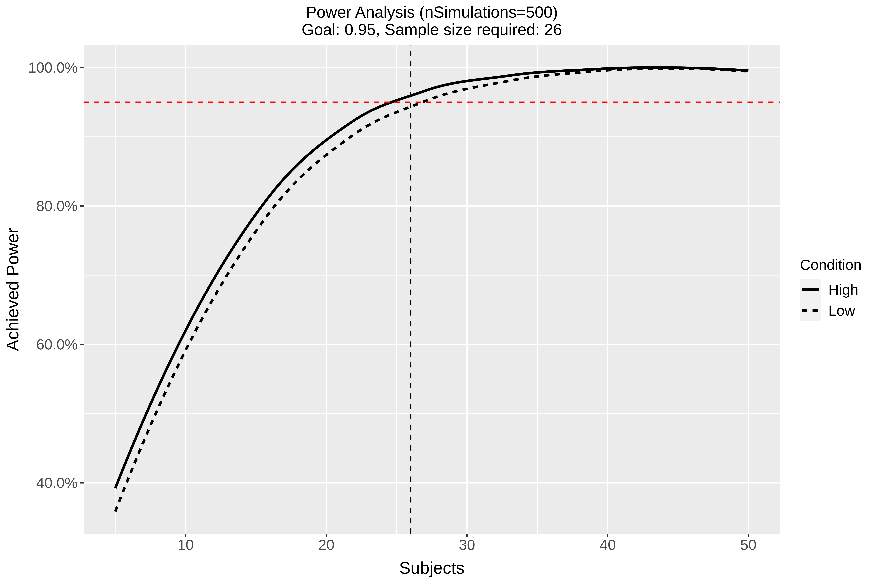

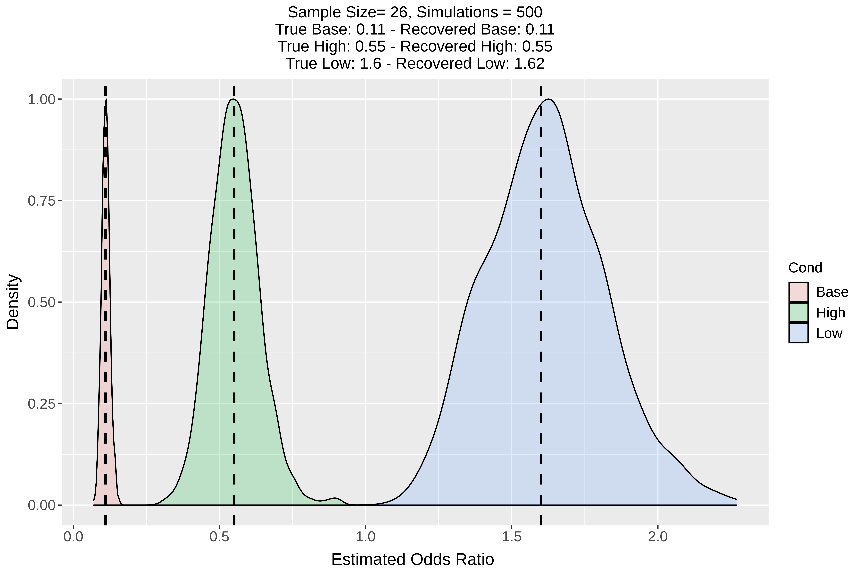

**b)**

**Supplementary Figure S2.** Simulation analysis for sample size calculation. a) We ran 500 simulations for sample sizes ranging from 5 to 50 participants to estimate the optimal sample size to achieve 95% power. Using a base odds ratio of .11 to report mind-blanking (MB) during free thinking, an odds ratio of 1.6 when arousal decreases ( *Low Arousal*, dotted line), and .55 when arousal increases (*High Arousal*, solid line), we estimated that a sample of 26 participants is sufficient to achieve significant power in both arousal conditions. b) To validate whether our model can recover the true parameters, we ran an additional 500 simulations using a sample size of 26 participants. Our results show that our model can indeed estimate the true parameters. Notes: dashed line = true parameters.

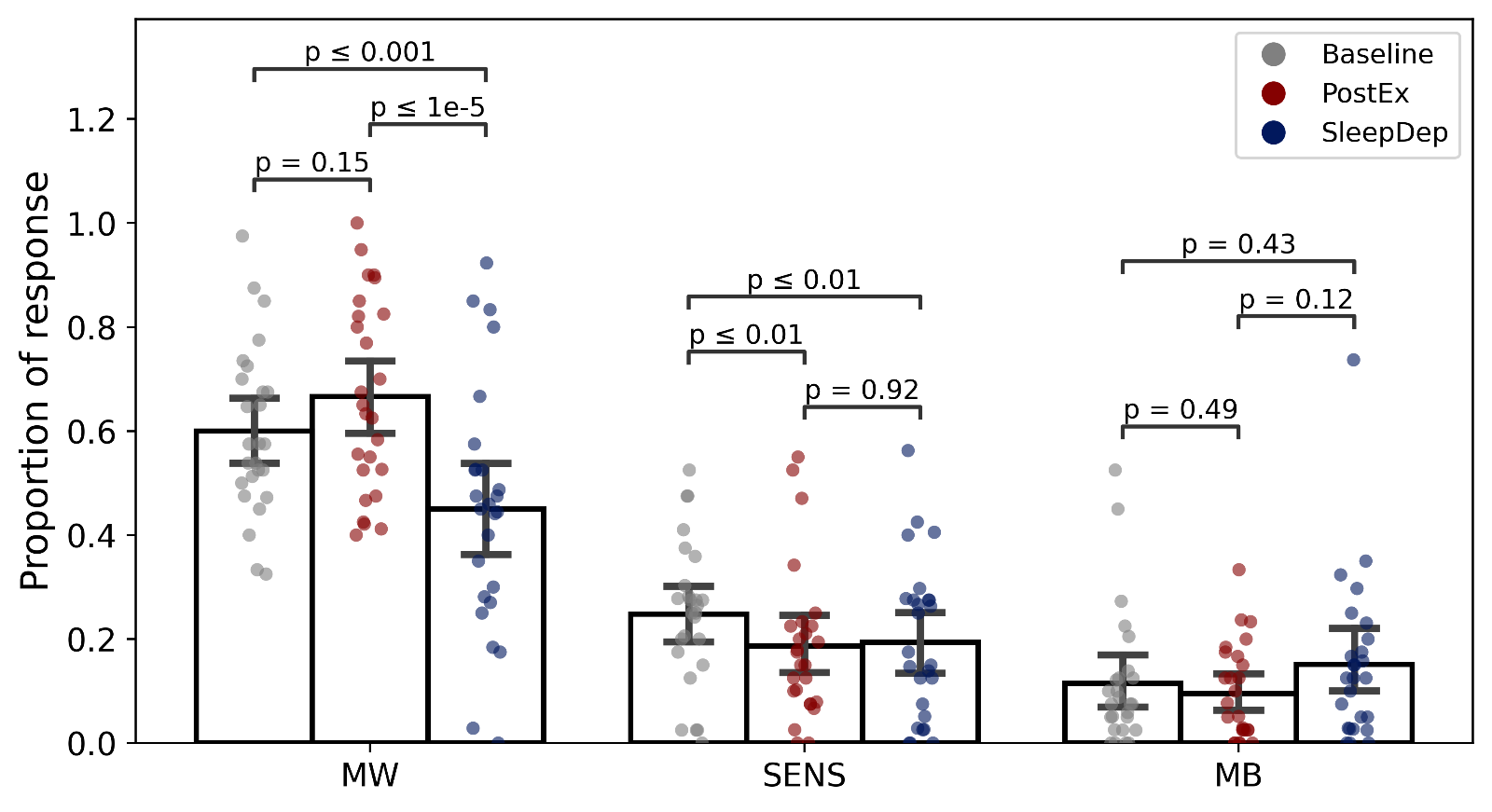

**Supplementary Figure S3.** General Estimating Equations (GEE) showed significant differences in mental state occurrences across arousal conditions. We found that while mind-wandering (MW) and sensations (SENS) reports decreased in *Low Arousal* compared to *Baseline* and *High Arousal*, there was no significant effect of *Low Arousal* on mind-blanking (MB) reports. Colored points indicate aggregate mean responses per subject. Errorbars indicate 95% confidence intervals.

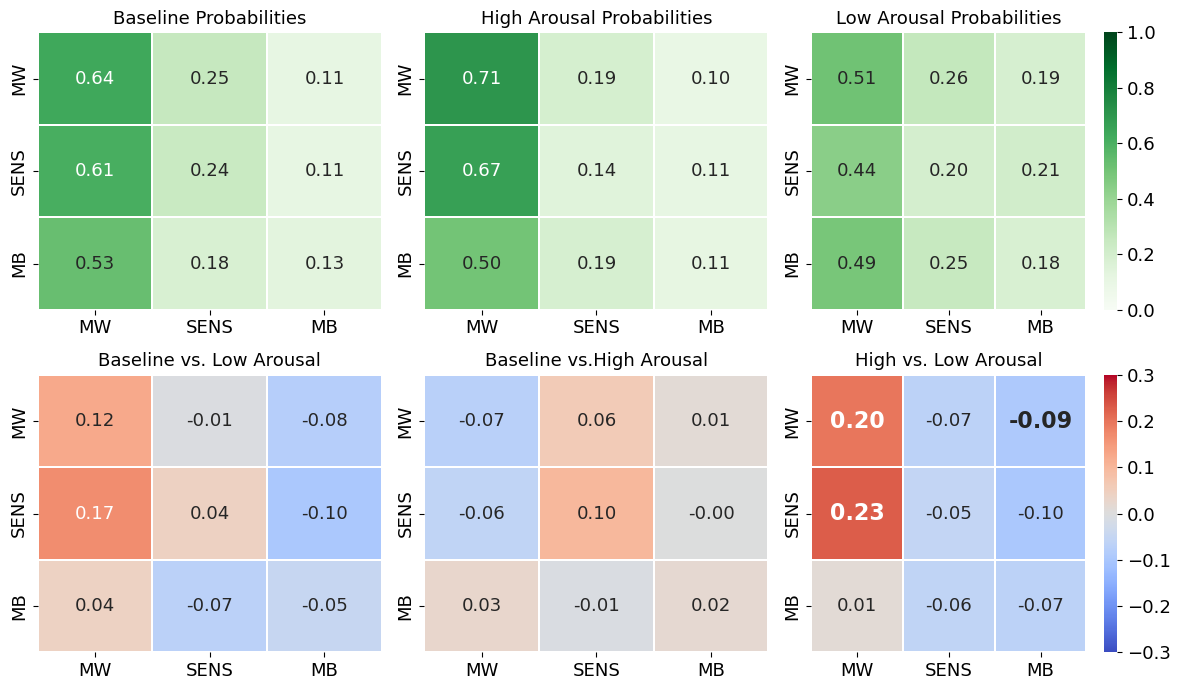

**Supplementary Figure S4.** Upper row: Mental state transitions varied across different arousal conditions. Average (green) transition probabilities across *Baseline*, *High* (after intense exercise) and *Low Arousal* (after sleep deprivation). Color bars and hue intensity indicate probability between 0-1. Lower row: Compared to *High Arousal*, participants were more likely to transition to MB and less likely to transition to MW in *Low Arousal*. Transition matrix difference across mental state reports. Numbers in bold indicate statistical significance (FDR corrected). The y-axis represents the origin (from), and the x-axis represents the direction (to).

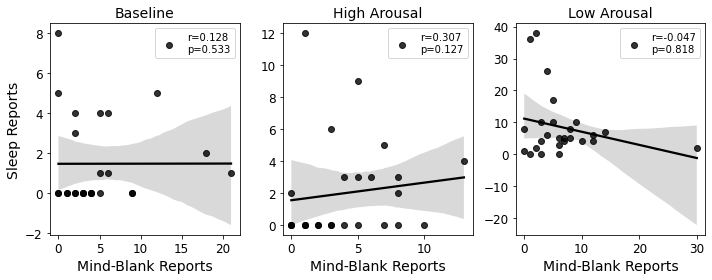

**Supplementary Figure S5*.*** MB reports did not correlate with Sleep reports across any arousal state. Lines indicate the best fit. Shaded areas represent 95% confidence intervals.

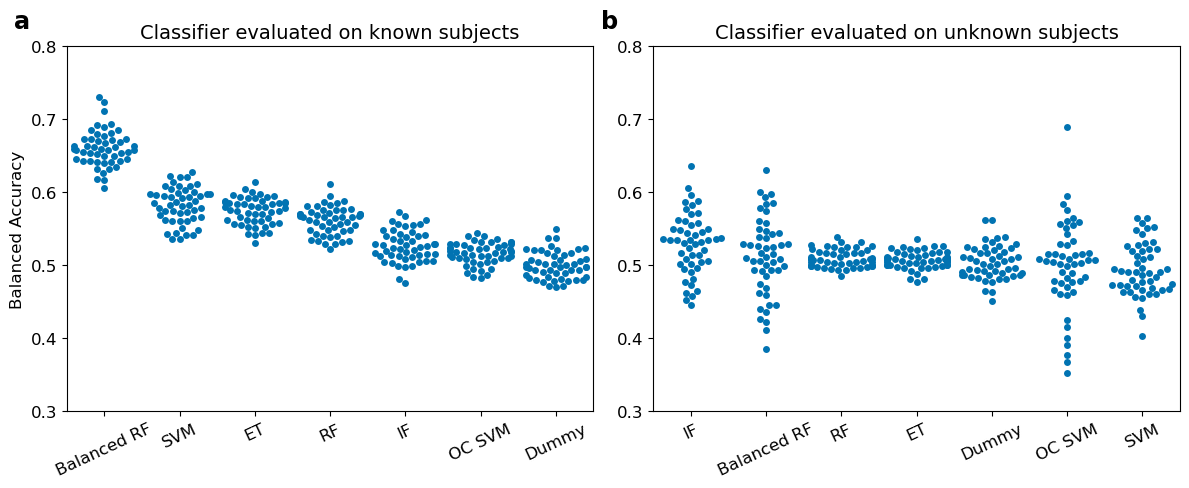

**Supplementary Figure S6.** Classification performance was above chance level when mind-blanking (MB) reports were pooled across subjects, but not after training on a subset of participants and classifying the remaining subset, when training the classifiers on the 1Hz filtered dataset. a) A balanced random forest classifier provided the highest classification performance across all examined classifiers including known subjects. b) An isolation forest classifier provided the highest classification performance across all examined classifiers on unknown samples. However, due to the high variance, we could not consider it meaningful. Individual points indicate performance on the folds of the repeated cross-validation. Results are ordered based on descending order of performance. Chance level performance is indicated by the Dummy classifier. RF = random forest; SVM = support vector machine; ET = extreme trees; IF = isolation forest; OC SVM = one-class support vector machine.

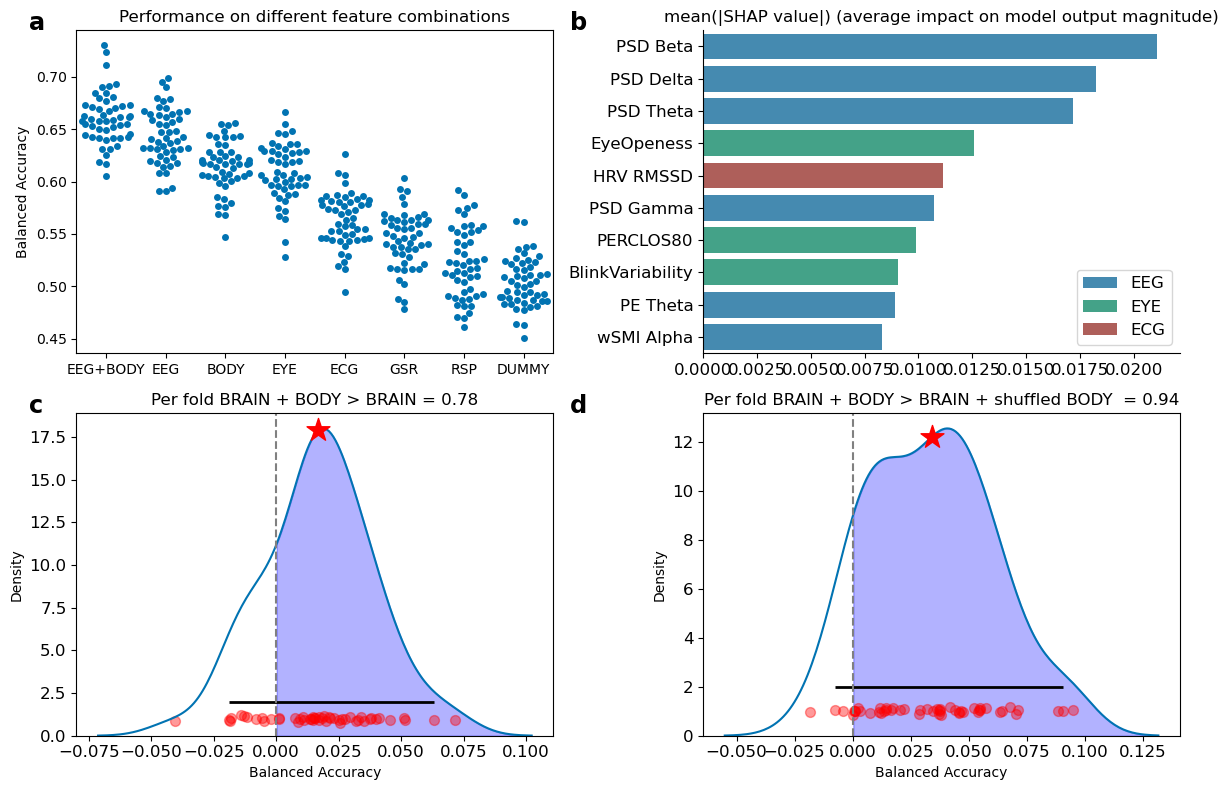

**Supplementary Figure S7.** a) A balanced random forest classifier trained on the 1Hz filtered dataset on a combination of BRAIN and BODY features outperformed classifiers trained solely on BRAIN or BODY features when evaluated with balanced accuracy. Individual points indicate performance on the folds of the repeated cross-validation. b) Subset of the 10 features with the highest mean of the absolute SHAP values obtained from the balanced random forest classifier. c) The per-fold differences between the classifier trained on both BRAIN and BODY features and the one trained only on BRAIN data suggest that incorporating both feature domains provides a slight performance improvement over using BRAIN data alone. The shaded region indicates better performance for the classifier trained on both feature domains. The star indicates the mean difference. The solid, horizontal line represents the 95% highest-density intervals of the distribution. Red dots indicate per-fold differences. d) The per-fold differences between the classifier trained on both BRAIN and BODY features and the one trained on BRAIN and shuffled BODY data suggest that the model with both BRAIN and BODY data does not consider the body markers as noise.

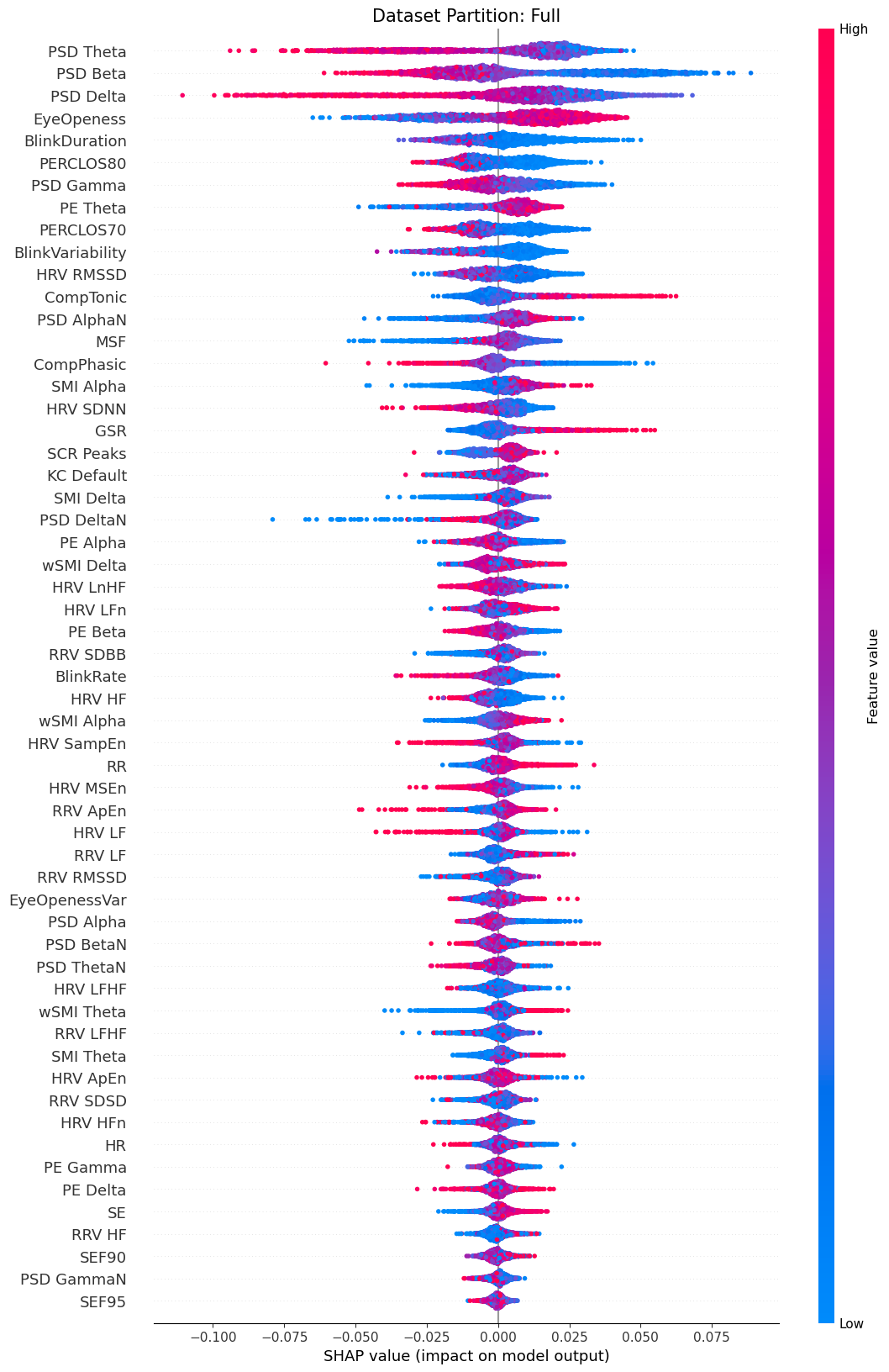

**Supplementary Figure S8.** SHAP values ranking of the .1 Hz filtered dataset. The SHAP value represents the impact of each feature on the model's prediction. Positive SHAP values push the prediction towards MB, while negative SHAP values push away from MB. Effectively, a high feature value with a high SHAP value indicates that when the feature increases, so does the probability of the model classifying a mental report as MB. Inversely, a high feature value with a low SHAP value indicates that when the feature increases, the probability of the model classifying a mental report as MB decreases.

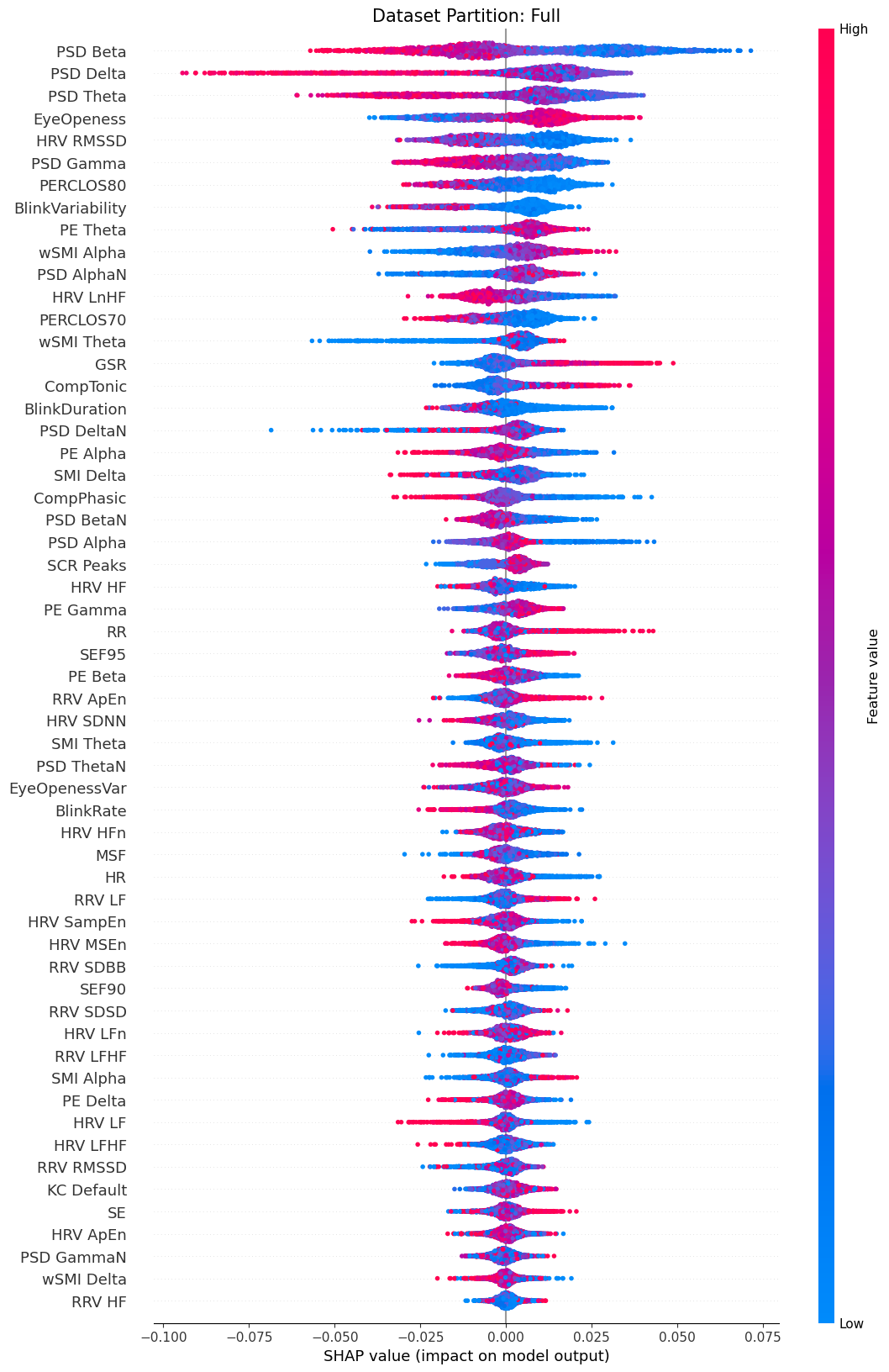

**Supplementary Figure S9.** SHAP values ranking of the 1 Hz filtered dataset. The SHAP value represents the impact of each feature on the model's prediction. Positive SHAP values push the prediction towards MB, while negative SHAP values push away from MB. Effectively, a high feature value with a high SHAP value indicates that when the feature increases, so does the probability of the model classifying a mental report as MB. Inversely, a high feature value with a low SHAP value indicates that when the feature increases, the probability of the model classifying a mental report as MB decreases.

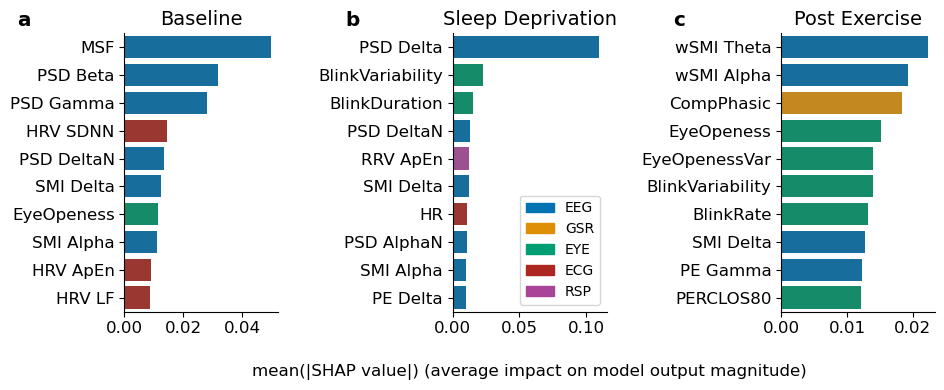

**Supplementary Figure S10.** Ranking of features by mean absolute SHAP value extracted from the balanced random forest classifier varied across different arousal conditions. a) Magnitude of SHAP values for a balanced random forest classifier trained on *Baseline Arousal* MB reports. The model relies mostly on features from the spectral domain of the EEG, the frequency domain of ECG, and eye openness. b) Magnitude for SHAP values for a classifier trained on *Low* *Arousal* MB reports. The model mostly uses spectral power in the delta band. c) Magnitude for SHAP values for a classifier trained on *High Arousal* MB reports. The model relies mostly on features from the connectivity in EEG, as well as EDA and eye openness.

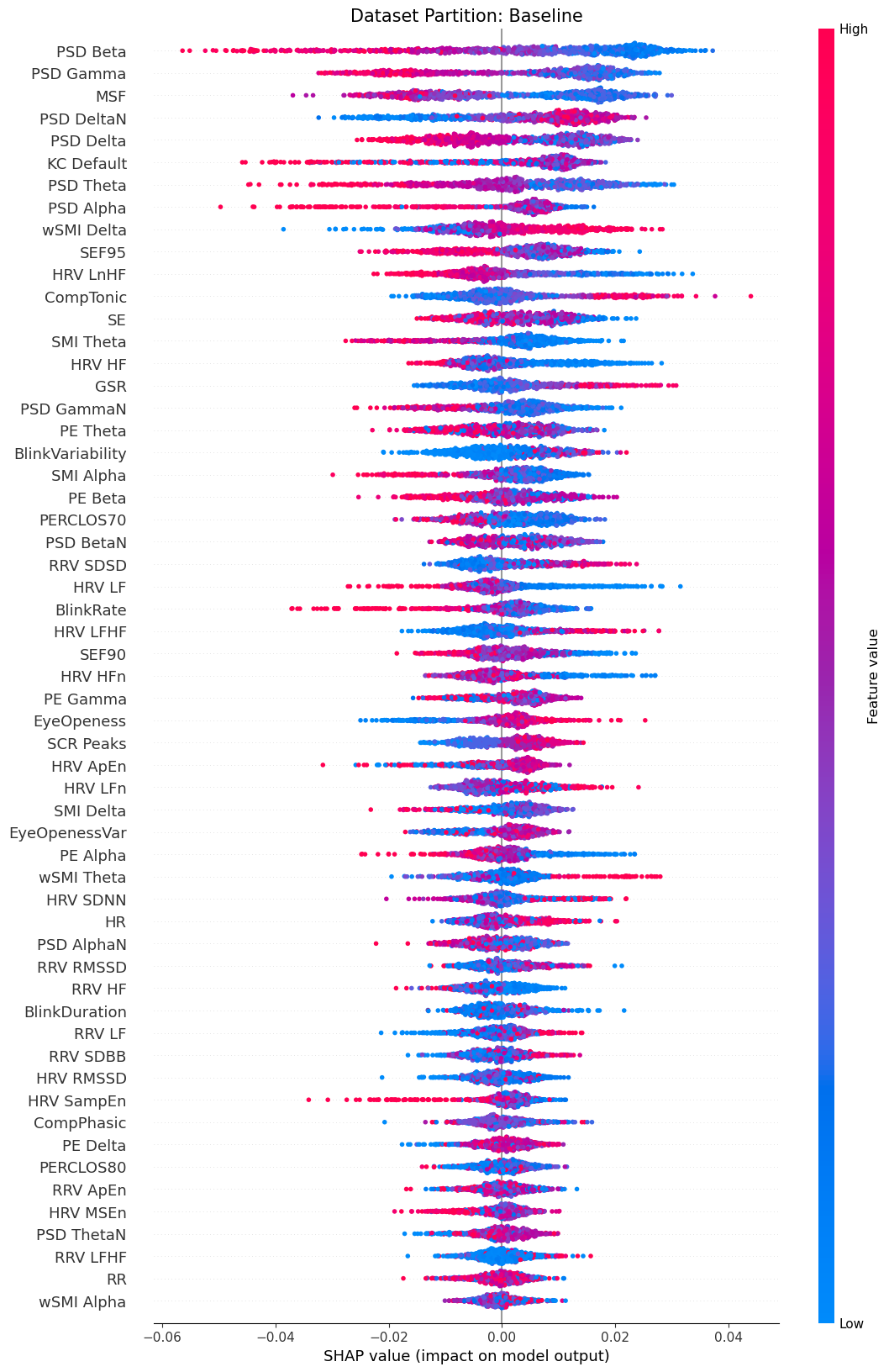

**Supplementary Figure S11.** SHAP values ranking for the *Baseline* mental state reports of the .1 Hz filtered dataset. The SHAP value represents the impact of each feature on the model's prediction. Positive SHAP values push the prediction towards MB, while negative SHAP values push away from MB. Effectively, a high feature value with a high SHAP value indicates that when the feature increases, so does the probability of the model classifying a mental report as MB. Inversely, a high feature value with a low SHAP value indicates that when the feature increases, the probability of the model classifying a mental report as MB decreases.

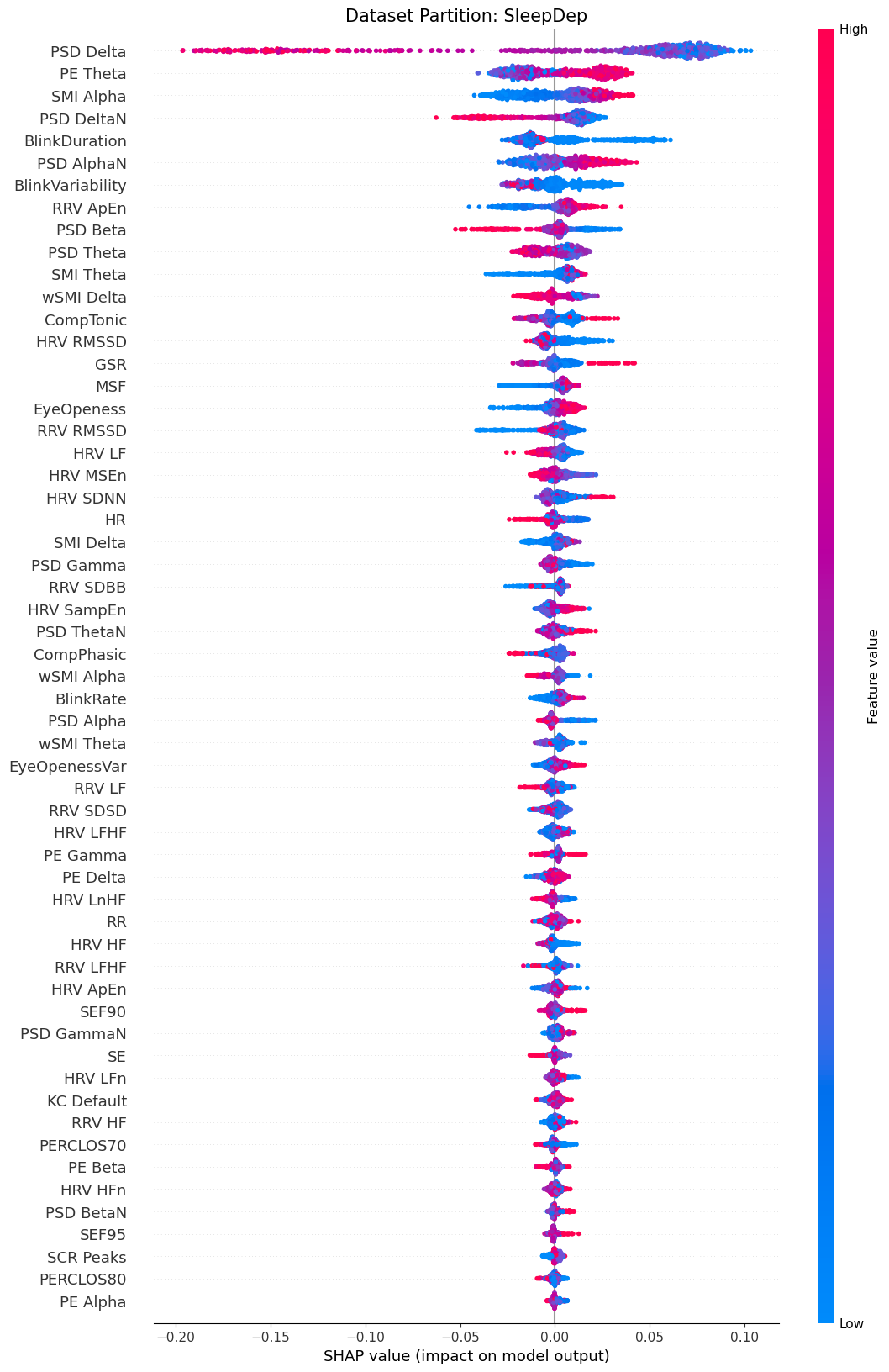

**Supplementary Figure S12.** SHAP values ranking for the *Low Arousal* mental state reports of the .1 Hz filtered dataset. The SHAP value represents the impact of each feature on the model's prediction. Positive SHAP values push the prediction towards MB, while negative SHAP values push away from MB. Effectively, a high feature value with a high SHAP value indicates that when the feature increases, so does the probability of the model classifying a mental report as MB. Inversely, a high feature value with a low SHAP value indicates that when the feature increases, the probability of the model classifying a mental report as MB decreases.

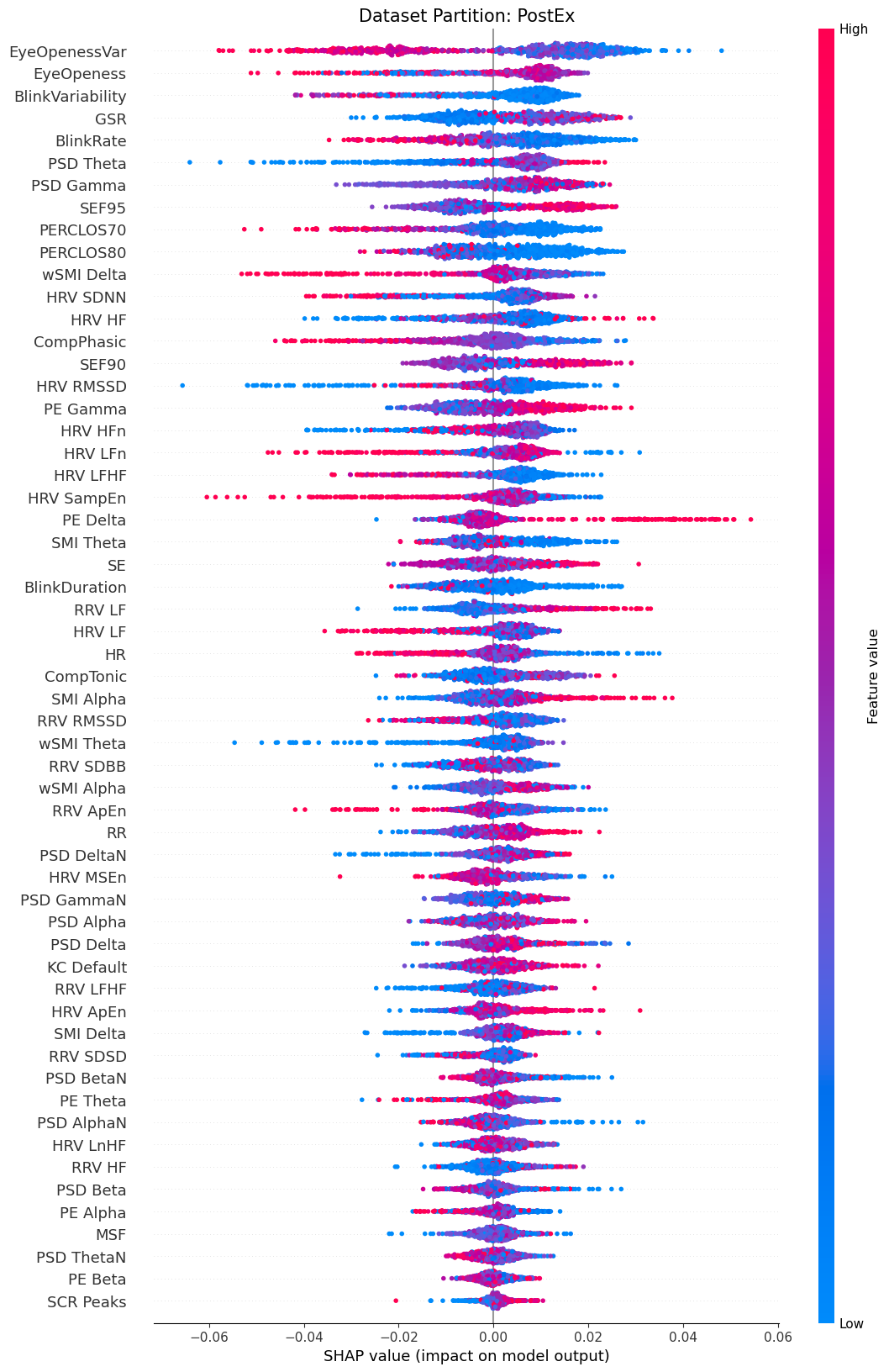

**Supplementary Figure S13.** SHAP values ranking for the *High Arousal* mental state reports of the .1 Hz filtered dataset. The SHAP value represents the impact of each feature on the model's prediction. Positive SHAP values push the prediction towards MB, while negative SHAP values push away from MB. Effectively, a high feature value with a high SHAP value indicates that when the feature increases, so does the probability of the model classifying a mental report as MB. Inversely, a high feature value with a low SHAP value indicates that when the feature increases, the probability of the model classifying a mental report as MB decreases.

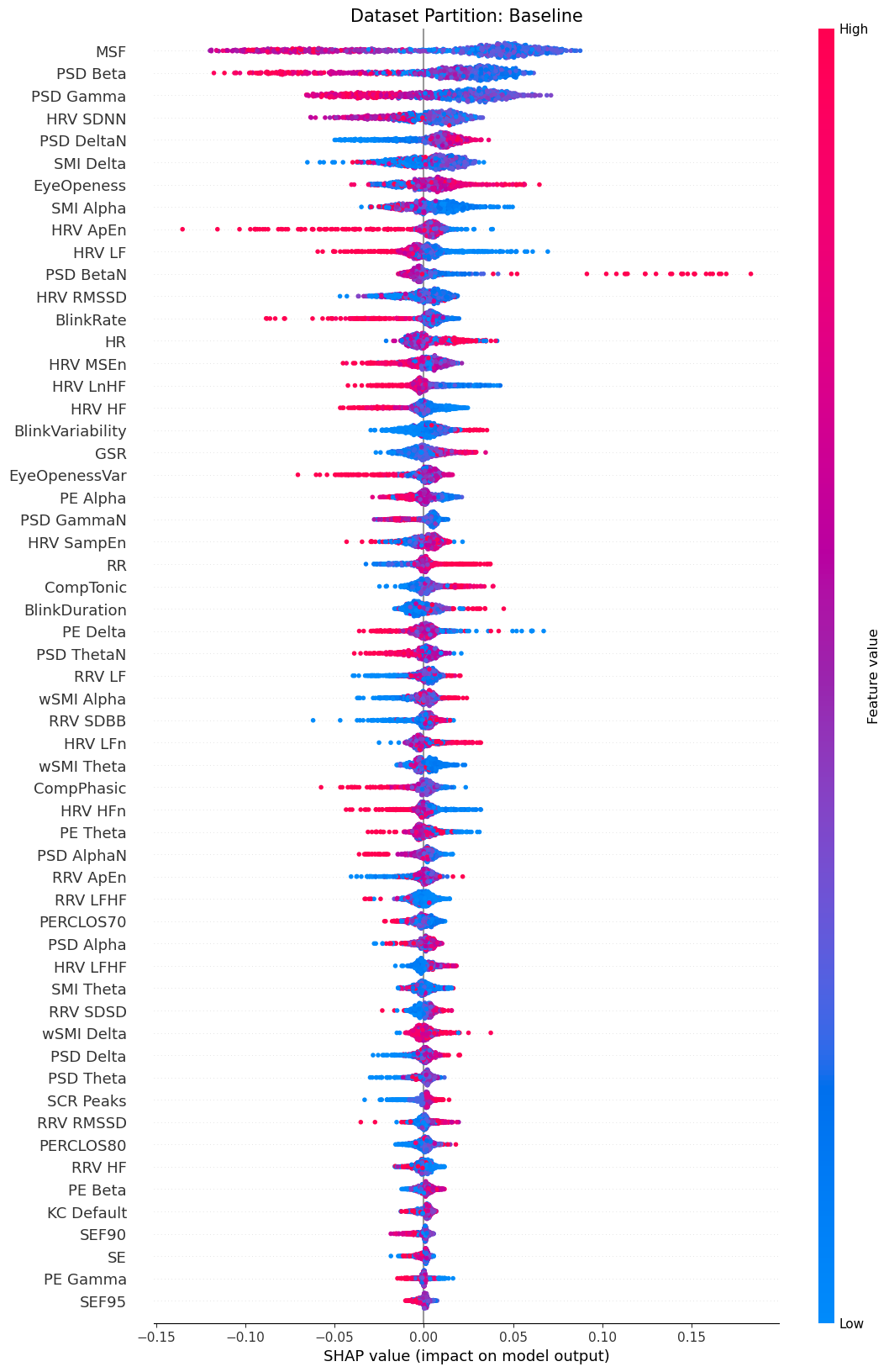

**Supplementary Figure S14.** SHAP values ranking for the *Baseline* mental state reports of the 1 Hz filtered dataset. The SHAP value represents the impact of each feature on the model's prediction. Positive SHAP values push the prediction towards MB, while negative SHAP values push away from MB. Effectively, a high feature value with a high SHAP value indicates that when the feature increases, so does the probability of the model classifying a mental report as MB. Inversely, a high feature value with a low SHAP value indicates that when the feature increases, the probability of the model classifying a mental report as MB decreases.

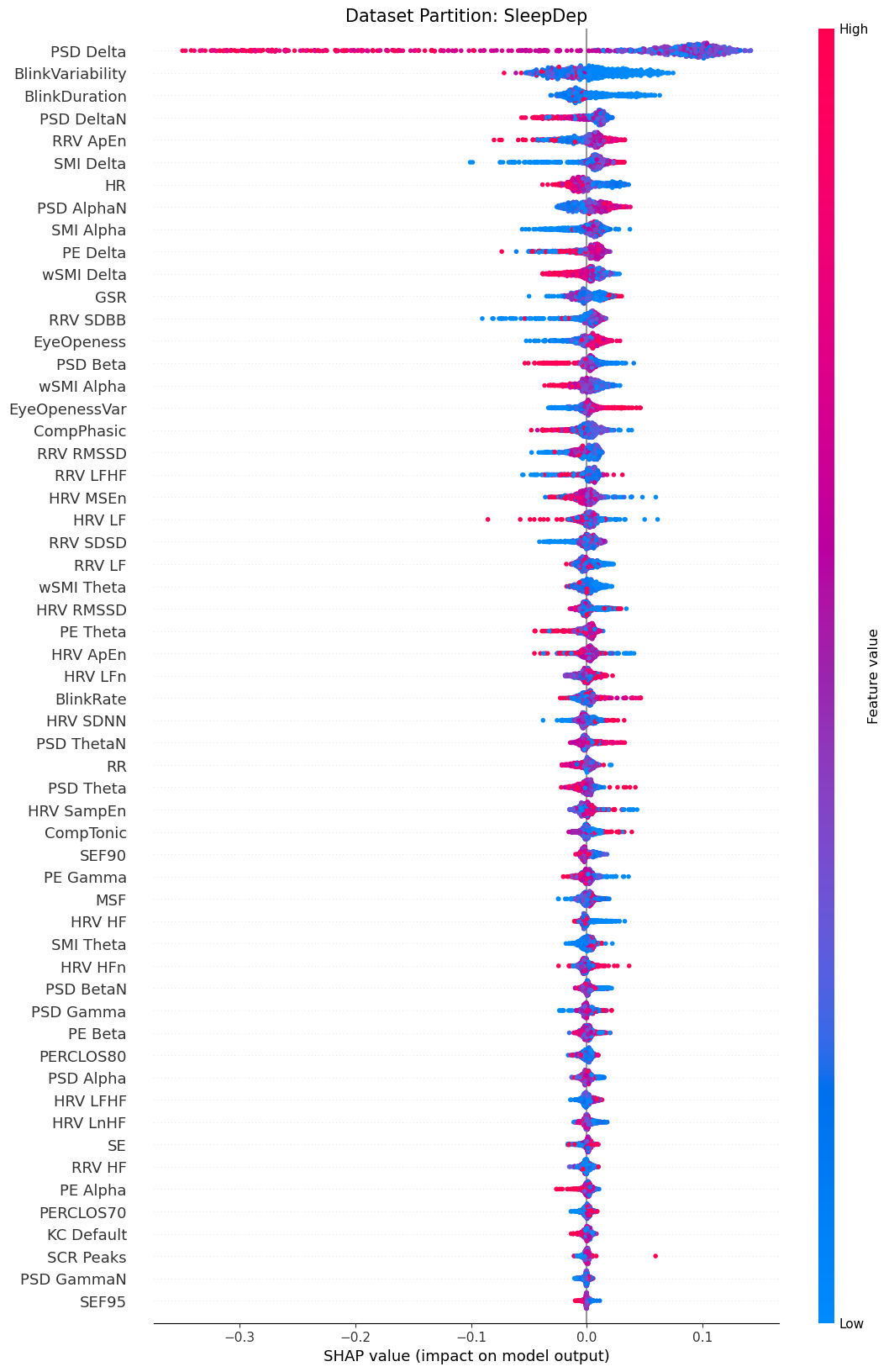

**Supplementary Figure S15.** SHAP values ranking for the *Low Arousal* mental state reports of the 1 Hz filtered dataset. The SHAP value represents the impact of each feature on the model's prediction. Positive SHAP values push the prediction towards MB, while negative SHAP values push away from MB. Effectively, a high feature value with a high SHAP value indicates that when the feature increases, so does the probability of the model classifying a mental report as MB. Inversely, a high feature value with a low SHAP value indicates that when the feature increases, the probability of the model classifying a mental report as MB decreases.

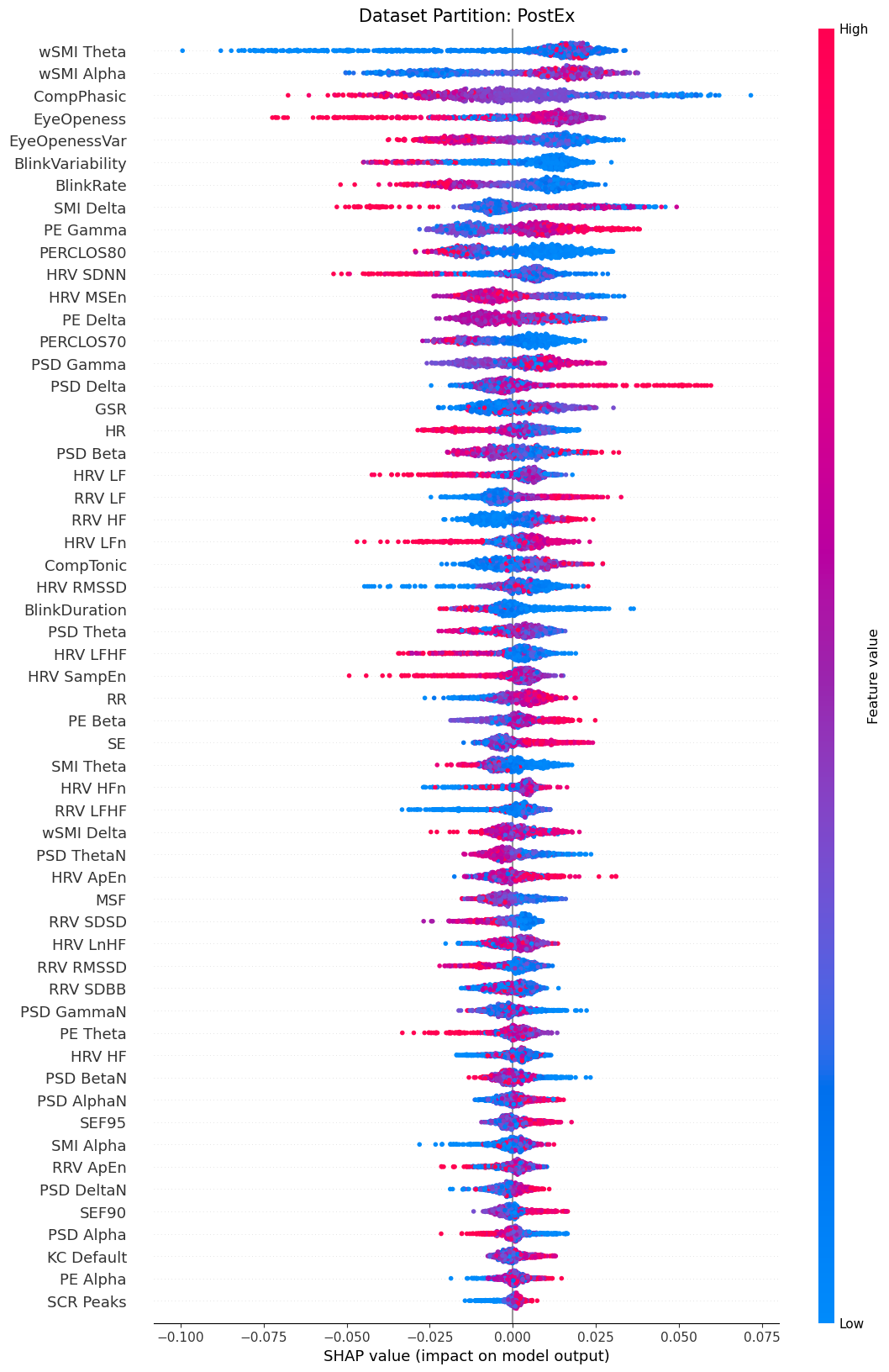

**Supplementary Figure S16.** SHAP values ranking for the *High Arousal* mental state reports of the 1 Hz filtered dataset. The SHAP value represents the impact of each feature on the model's prediction. Positive SHAP values push the prediction towards MB, while negative SHAP values push away from MB. Effectively, a high feature value with a high SHAP value indicates that when the feature increases, so does the probability of the model classifying a mental report as MB. Inversely, a high feature value with a low SHAP value indicates that when the feature increases, the probability of the model classifying a mental report as MB decreases.

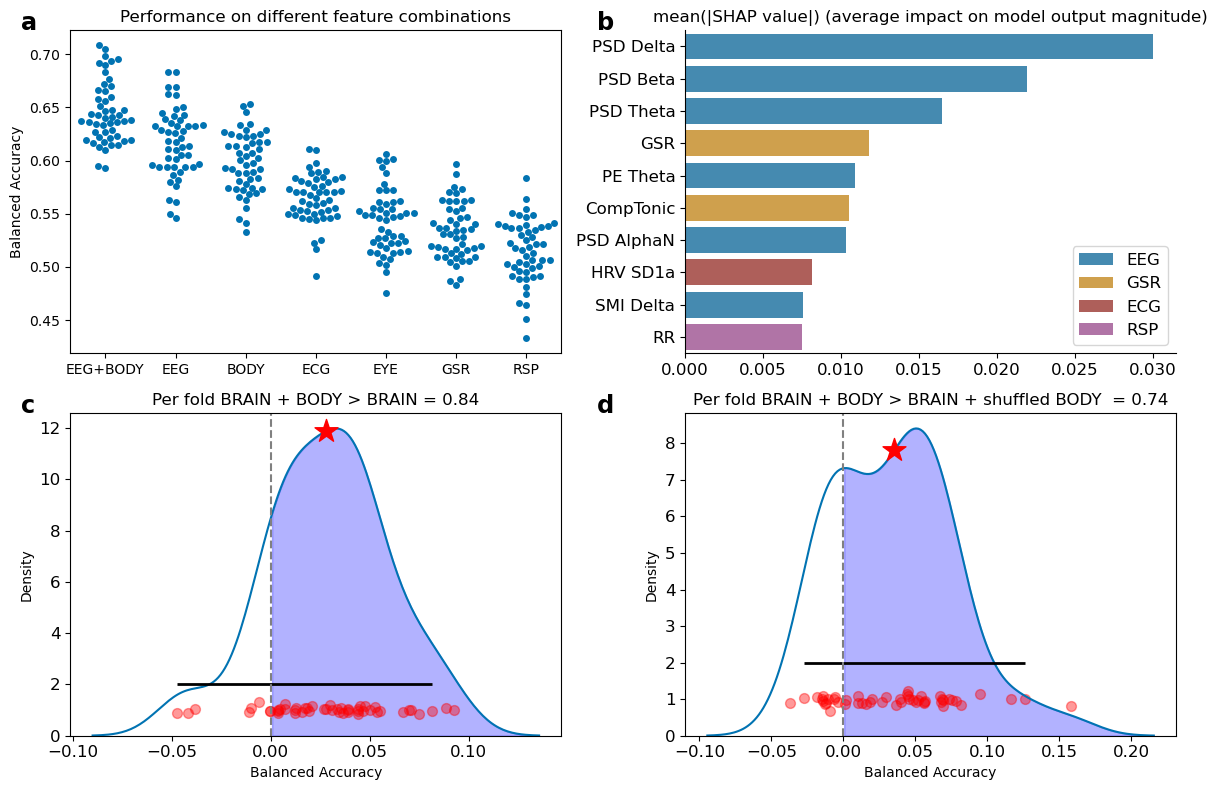

**Supplementary Figure S17.** a) A balanced random forest classifier trained on the last 10 seconds of the .1Hz filtered dataset on a combination of BRAIN and BODY features outperformed classifiers trained solely on BRAIN or BODY features when evaluated with balanced accuracy. Individual points indicate performance on the folds of the repeated cross-validation. b) Subset of the 10 features with the highest mean of the absolute SHAP values obtained from the balanced random forest classifier. c) The per-fold differences between the classifier trained on both BRAIN and BODY features and the one trained only on BRAIN data suggest that incorporating both feature domains provides a slight performance improvement over using BRAIN data alone. The shaded region indicates better performance for the classifier trained on both feature domains. The star indicates the mean difference. The solid, horizontal line represents the 95% highest-density intervals of the distribution. Red dots indicate per-fold differences. d) The per-fold differences between the classifier trained on both BRAIN and BODY features and the one trained on BRAIN and shuffled BODY data suggest that the model with both BRAIN and BODY data does not consider the body markers as noise.

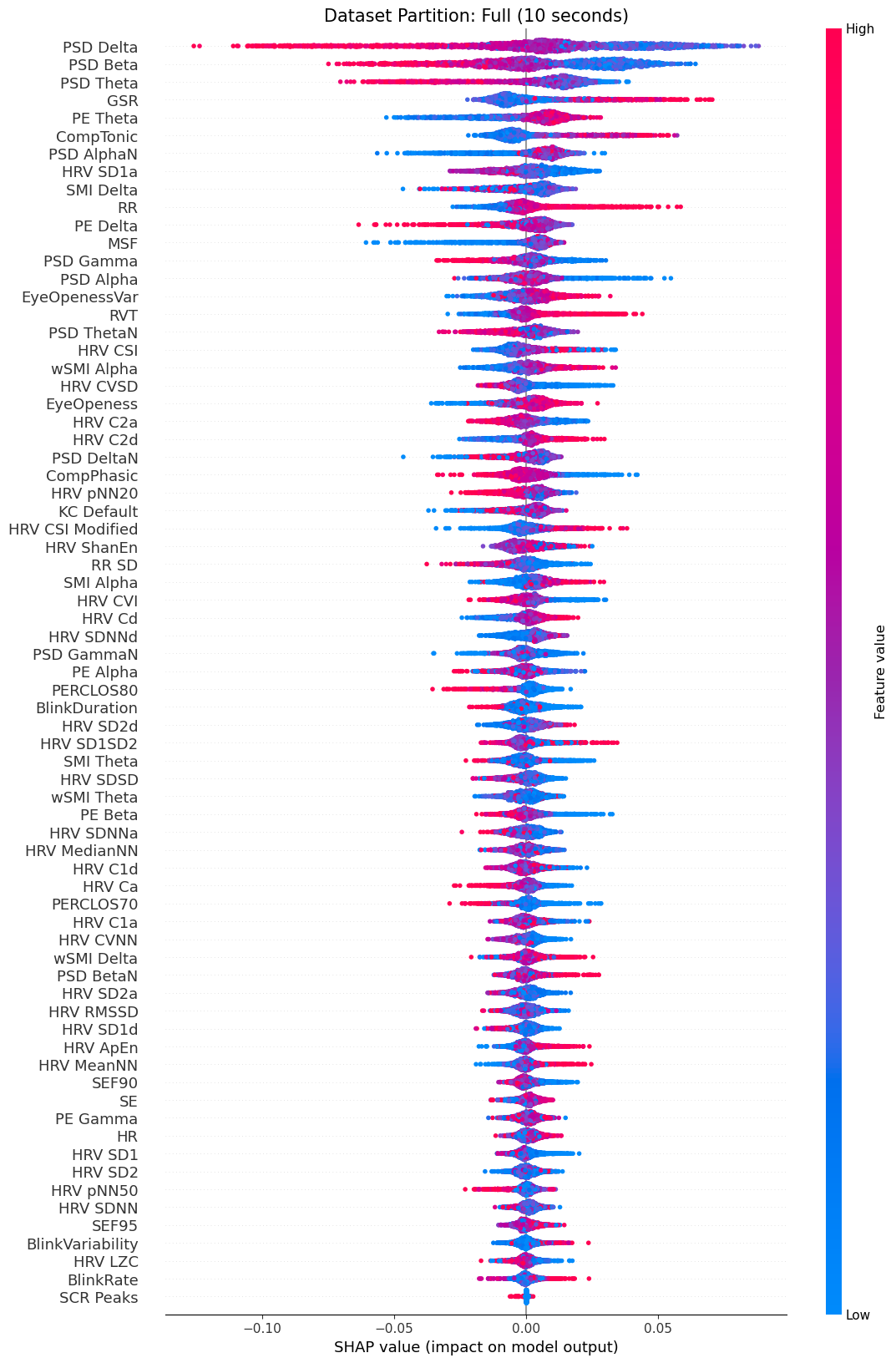

**Supplementary Figure S18.** SHAP values ranking of the .1Hz filtered dataset (last 10 seconds partition). The SHAP value represents the impact of each feature on the model's prediction. Positive SHAP values push the prediction towards MB, while negative SHAP values push away from MB. Effectively, a high feature value with a high SHAP value indicates that when the feature increases, so does the probability of the model classifying a mental report as MB. Inversely, a high feature value with a low SHAP value indicates that when the feature increases, the probability of the model classifying a mental report as MB decreases.

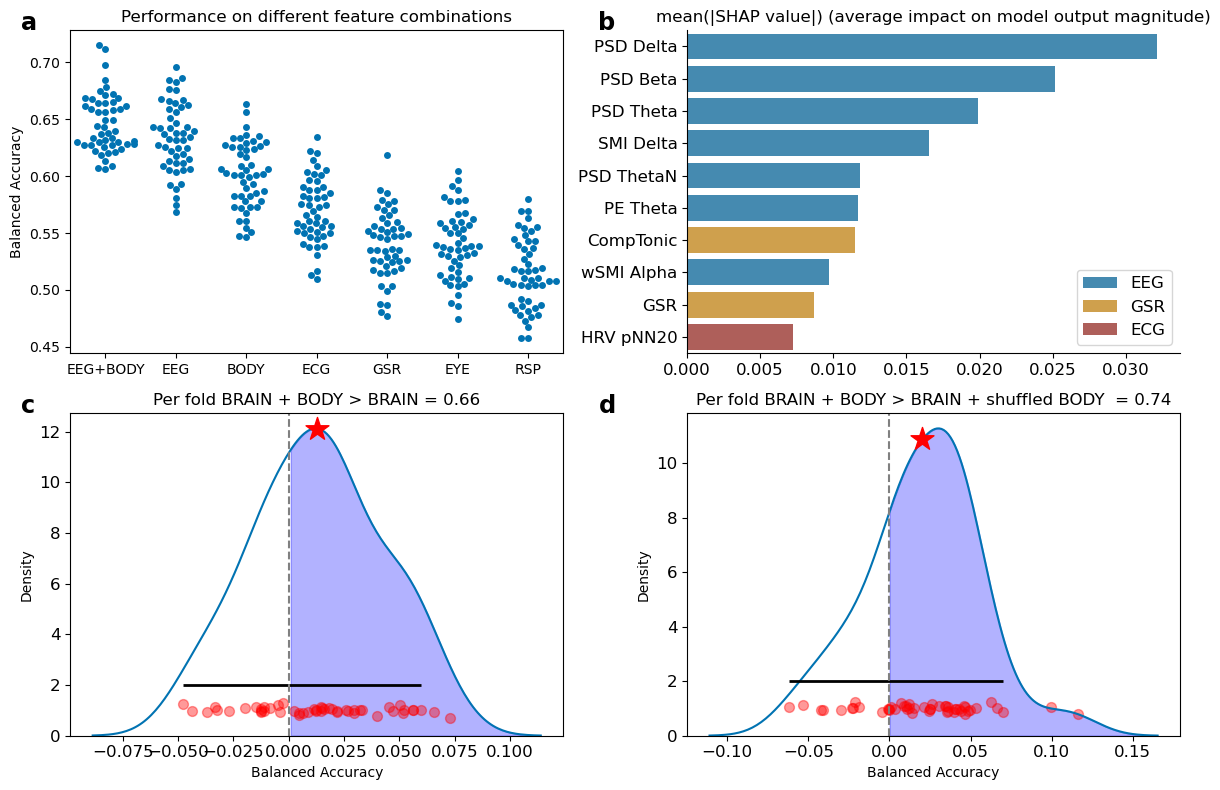

**Supplementary Figure S19.** a) A balanced random forest classifier trained on the last 10 seconds of the 1Hz filtered dataset on a combination of BRAIN and BODY features outperformed classifiers trained solely on BRAIN or BODY features when evaluated with balanced accuracy. Individual points indicate performance on the folds of the repeated cross-validation. b) Subset of the 10 features with the highest mean of the absolute SHAP values obtained from the balanced random forest classifier. c) The per-fold differences between the classifier trained on both BRAIN and BODY features and the one trained only on BRAIN data suggest that incorporating both feature domains provides a slight performance improvement over using BRAIN data alone. The shaded region indicates better performance for the classifier trained on both feature domains. The star indicates the mean difference. The solid, horizontal line represents the 95% highest-density intervals of the distribution. Red dots indicate per-fold differences. d) The per-fold differences between the classifier trained on both BRAIN and BODY features and the one trained on BRAIN and shuffled BODY data suggest that the model with both BRAIN and BODY data does not consider the body markers as noise.

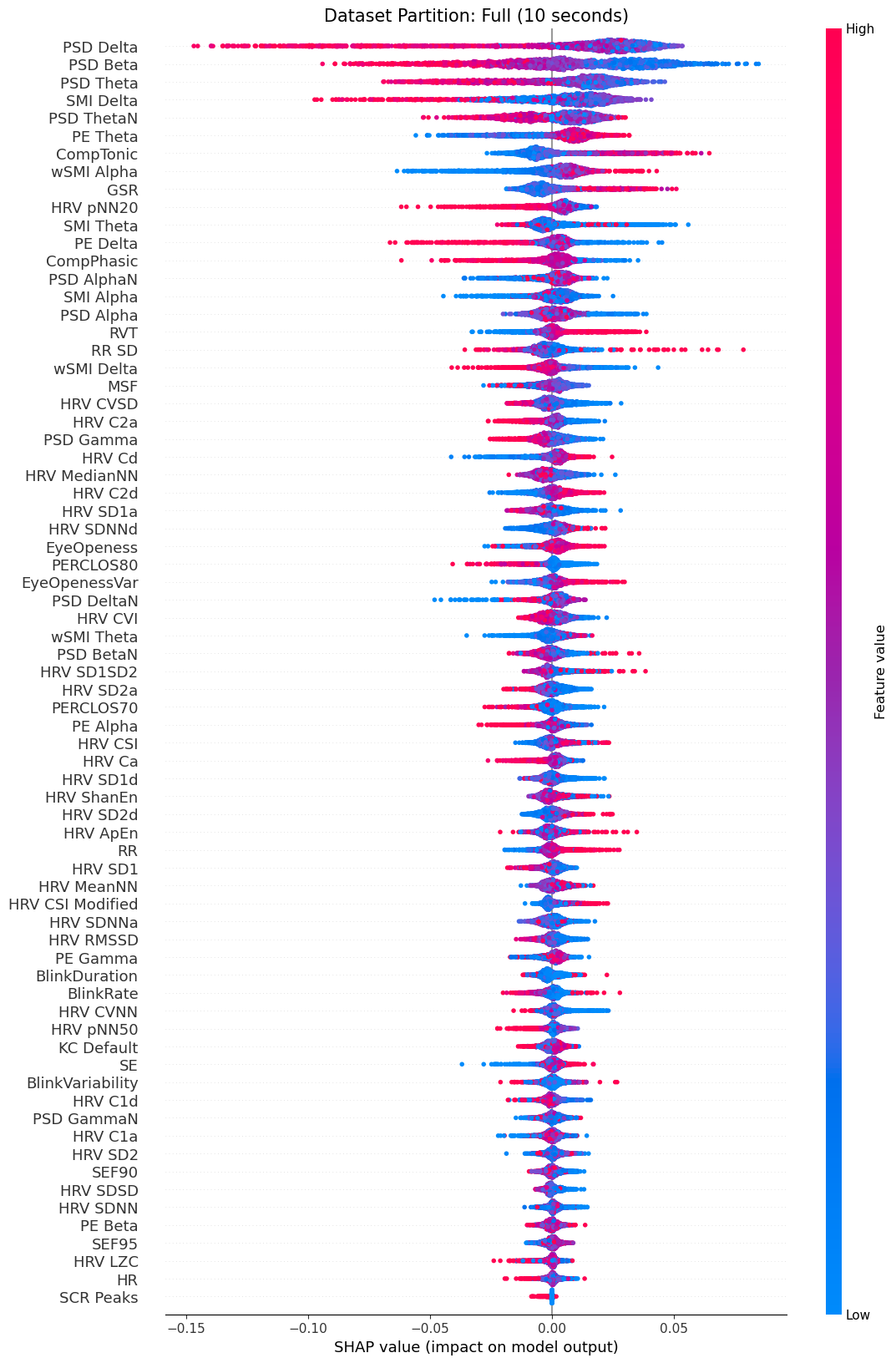

**Supplementary Figure S20.** SHAP values ranking of the 1Hz filtered dataset (last 10 seconds partition). The SHAP value represents the impact of each feature on the model's prediction. Positive SHAP values push the prediction towards MB, while negative SHAP values push away from MB. Effectively, a high feature value with a high SHAP value indicates that when the feature increases, so does the probability of the model classifying a mental report as MB. Inversely, a high feature value with a low SHAP value indicates that when the feature increases, the probability of the model classifying a mental report as MB decreases.

| **Supplementary Table S1**. Design Table | | | | |
| --- | --- | --- | --- | --- |
| **Question** | **Hypothesis** | **Sampling Plan** | **Analysis Plan** | **Alternative Explanation** |
| Is automimic arousal implicated in mental state reportability? | Low and high arousal promote more frequent MB reports. | 500 Simulations for datasets ranging from 5 to 50 participants.  For each dataset, we fit a binomial model with an odds ratio of .1 for MB occurrence  N=26 participants | MB ~ Arousal  Mental Report ~ Arousal  RT ~ Mental Report  Transitions ~ Mental Report | Low arousal manipulation was not effective in modifying autonomic signals.  High arousal manipulation did not last throughout the experience-sampling procedure.  Higher arousal levels might facilitate monitoring, reducing MB reports. |
| Can mental absences be attributed to cerebral mechanisms only, or to brain-body interactions? | We can decode MB from other mental reports based on a brain-body profile characterized by lower overall complexity. | NA | Train 4 classifiers:   - 1.Support Vector Machine - 2.One class SVM - 3.Random Forest - 4. Random Trees   Optimize for F1-score  Nested CV hyperparameter tuning | Given the unbalanced nature of our dataset, our classifiers might not converge properly to accurate prediction parameters.  Physiological timeseries might be too slow (few oscillations) to contribute to short events, such as a mental state. |

| **Supplementary Table S2**. A brief overview of the effects of sleep deprivation and exercise on arousal metrics | | |
| --- | --- | --- |
| Modality | Metric | Previous Studies |
| Electrocardiogram (EEG) | Alpha oscillations  Delta oscillations  Theta oscillations | [1,3] |
| Electroencephalogram (ECG) | Heart Rate  Heart Rate Variability | [4, 5] |
| Pupillometry | Pupil Size | [6, 7] |

| **Supplementary Table S3.** People tended to give reports faster in *Baseline Arousal.* Additionally, people tend to report mind-wandering (MW) the fastest. The bold font indicates significance. | | | | |
| --- | --- | --- | --- | --- |
| **Contrast** | **Marginal Mean** | **SE** | **CL** | **pFDR** |
| Baseline - High  Baseline - Low  High-Low  MB - MW  MB - SENS  MW-SENS  MW Baseline – MW High  MW Baseline – MW Low  MW High – MW Low  SENS Baseline – SENS High  SENS Baseline – SENS Low  SENS High – SENS Low  MB Baseline – MB High  MB Baseline – MB Low  MB High – MB Low | .02 | .01 | [.00,.03] | **7.3e-04** |
|  | .03 | .00 | [.02,.04] | **5e-09** |
|  | .01 | .00 | [0,.03] | **1.1e-02** |
|  | -.09 | .00 | [ -.09, -.07] | **1.5e-59** |
|  | -.09 | .01 | [ -.1, -.07] | **2.6e-50** |
|  | -.01 | .00 | [ -.02, .01] | 1.2e-01 |
|  | .01 | .00 | [.00, .02] | **1.1e-02** |
|  | .06 | .00 | [.04, .08] | **1.1e-21** |
|  | .05 | .01 | [.04, .07] | 2e-19 |
|  | .02 | .01 | [-.01, .05] | 1.7e-01 |
|  | .01 | .01 | [-.02, .04] | 4.1e-01 |
|  | -.01 | .01 | [-.04, .02] | 4.5e-01 |
|  | .03 | .01 | [.00, .06] | **1.2e-02** |
|  | .02 | .01 | [-.01, .05] | **3.4e-02** |
|  | -.01 | .01 | [-.04, .02] | 4.3e-01 |

| **Supplementary Table S4.** A balanced random forest classifier trained on the 1Hz filtered dataset outperformed all classifiers when compared across balanced accuracy. Cells indicate mean and 95% CI. | | | | | | |
| --- | --- | --- | --- | --- | --- | --- |
| **Examined** | **Classifier** | **Recall** | **Precision** | **F1** | **ROC AUC** | **Balanced Accuracy** |
| Known Subjects | Balanced RF | .61, [.6, .63] | .26, [.26, .27] | .37, [.36, .37] | .71, [.7, .72] | .66, [.65, .67] |
|  | SVM | .28, [.26, .3] | .29, [.28, .31] | .28, [.27, .3] | .63, [.62, .64] | .58, [.58, .59] |
|  | ET | .16, [.15, .17] | .64, [.6, .67] | .26, [.24, .27] | .72, [.71, .73] | .57, [.57, .58] |
|  | RF | .14, [.13, .15] | .61, [.57, .64] | .22, [.21, .24] | .7, [.69, .71] | .56, [.56, .57] |
|  | IF | .15, [.14, .16] | .21, [.19, .22] | .17, [.16, .18] | .53, [.52, .53] | .53, [.52, .53] |
|  | OC SVM | .91, [.89, .93] | .15, [.15, .15] | .26, [.25, .26] | .52, [.51, .52] | .52, [.51, .52] |
|  | DUMMY | .14, [.13, .15] | .15, [.14, .15] | .14, [.13, .15] | .5, [.5, .51] | .5, [.49, .51] |
| Unknown Subjects | IF | .25, [.21, .29] | .18, [.16, .2] | .2, [.18, .22] | .53, [.52, .54] | .53, [.52, .54] |
|  | RF | .04, [.03, .05] | .27, [.21, .34] | .06, [.05, .08] | .53, [.51, .54] | .51, [.51, .51] |
|  | Balanced RF | .37, [.32, .42] | .16, [.14, .18] | .21, [.18, .23] | .51, [.49, .53] | .51, [.5, .53] |
|  | ET | .03, [.02, .03] | .34, [.26, .43] | .05, [.04, .06] | .51, [.5, .53] | .51, [.5, .51] |
|  | DUMMY | .15, [.14, .17] | .15, [.13, .17] | .15, [.14, .16] | .5, [.49, .5] | .5, [.5, .51] |
|  | OC SVM | .69, [.61, .77] | .15, [.13, .16] | .23, [.21, .24] | .5, [.48, .52] | .5, [.48, .52] |
|  | SVM | .2, [.18, .22] | .14, [.13, .16] | .16, [.14, .17] | .48, [.47, .5] | .49, [.48, .5] |
| RF = Random Forest; SVM = Support Vector Machine; ET = Extreme Trees; IF = Isolation Forest;  OC SVM = One-Class Support Vector Machine | | | | | | |

| **Supplementary Table S5**. A classifier trained on a combination of BRAIN and BODY features outperformed classifiers trained solely on BRAIN or BODY features on the 1Hz filtered dataset when evaluated with balanced accuracy. Cells indicate mean and 95% CI. | | | | | |
| --- | --- | --- | --- | --- | --- |
| **Classifier** | **Recall** | **Precision** | **F1** | **ROC AUC** | **Balanced Accuracy** |
| BRAIN + BODY | .61, [.6, .63] | .26, [.26, .27] | .37, [.36, .37] | .71, [.7, .72] | .66, [.65, .67] |
| BRAIN | .58, [.56, .6] | .25, [.25, .26] | .35, [.34, .36] | .69, [.68, .7] | .64, [.64, .65] |
| BODY | .6, [.58, .61] | .22, [.21, .22] | .32, [.31, .32] | .67, [.66, .68] | .61, [.61, .62] |
| EYE | .57, [.55, .58] | .22, [.21, .22] | .31, [.3, .32] | .64, [.63, .65] | .61, [.6, .62] |
| ECG | .56, [.54, .57] | .18, [.17, .18] | .27, [.26, .28] | .59, [.58, .6] | .56, [.56, .57] |
| EDA | .6, [.57, .63] | .17, [.16, .17] | .26, [.25, .26] | .57, [.56, .58] | .54, [.54, .55] |
| RSP | .54, [.52, .55] | .15, [.15, .16] | .24, [.23, .25] | .53, [.52, .54] | .52, [.51, .53] |

| **Supplementary Table S6**. Classification performance was retained when examining mind-blanking (MB) occurrences from each arousal condition separately on the .1 Hz filtered dataset. Cells indicate mean and 95% CI. | | | | | |
| --- | --- | --- | --- | --- | --- |
| **Arousal** | **Recall** | **Precision** | **F1** | **ROC AUC** | **Balanced Accuracy** |
| Baseline | .62, [.59, .65] | .24, [.23, .25] | .34, [.33, .36] | .73, [.71, .74] | .67, [.65, .68] |
| Low | .57, [.54, .6] | .34, [.33, .35] | .42, [.41, .43] | .7, [.69, .71] | .64, [.63, .65] |
| High | .61, [.58, .64] | .16, [.16, .17] | .26, [.25, .27] | .66, [.64, .68] | .61, [.6, .63] |

| **Supplementary Table S7**. Classification performance was retained when examining mind-blanking (MB) occurrences from each arousal condition separately on the 1 Hz filtered dataset. Cells indicate mean and 95% CI. | | | | | |
| --- | --- | --- | --- | --- | --- |
| **Arousal** | **Recall** | **Precision** | **F1** | **ROC AUC** | **Balanced Accuracy** |
| Baseline | .62, [.59, .64] | .24, [.23, .24] | .34, [.33, .35] | .72, [.7, .74] | .66, [.65, .68] |
| Low | .59, [.56, .61] | .34, [.33, .35] | .43, [.42, .44] | .71, [.7, .72] | .65, [.63, .66] |
| High | .58, [.55, .62] | .16, [.15, .17] | .25, [.24, .26] | .65, [.64, .67] | .6, [.59, .62] |

| **Supplementary Table S8.** Classifier performance on different feature combinations using 10s of data, filtered at .1 Hz. Performance did not alter when using the BRAIN markers but was reduced in the BODY markers. Cells indicate mean and 95% CI. | | | | | |
| --- | --- | --- | --- | --- | --- |
| **Classifier** | **Recall** | **Precision** | **F1** | **ROC AUC** | **Balanced Accuracy** |
| EEG + BODY | .62, [.6, .63] | .24, [.23, .24] | .34, [.34, .35] | .69, [.68, .7] | .65, [.64, .65] |
| EEG | .58, [.56, .6] | .22, [.21, .22] | .32, [.31, .32] | .66, [.65, .67] | .62, [.61, .63] |
| BODY | .58, [.57, .6] | .2, [.19, .2] | .3, [.29, .3] | .64, [.63, .64] | .6, [.59, .61] |
| ECG | .52, [.5, .53] | .18, [.18, .18] | .27, [.26, .27] | .59, [.58, .59] | .56, [.56, .57] |
| EDA | .51, [.49, .54] | .17, [.16, .17] | .25, [.24, .26] | .56, [.55, .57] | .54, [.53, .55] |
| EYE | .64, [.61, .67] | .16, [.15, .16] | .25, [.24, .26] | .56, [.55, .57] | .53, [.53, .54] |
| RSP | .67, [.63, .71] | .15, [.14, .15] | .24, [.23, .25] | .52, [.51, .53] | .51, [.51, .52] |

| **Supplementary Table S9.** Classifier performance on different feature combinations using 10s of data, filtered at 1 Hz. Performance did not alter when using the BRAIN markers but was reduced in the BODY markers. Cells indicate mean and 95% CI. | | | | | |
| --- | --- | --- | --- | --- | --- |
| **Classifier** | **Recall** | **Precision** | **F1** | **ROC AUC** | **Balanced Accuracy** |
| EEG + BODY | .62, [.6, .63] | .24, [.24, .25] | .35, [.34, .35] | .69, [.69, .7] | .65, [.64, .65] |
| EEG | .59, [.58, .61] | .23, [.23, .24] | .33, [.33, .34] | .68, [.67, .69] | .63, [.63, .64] |
| BODY | .58, [.57, .6] | .2, [.2, .21] | .3, [.29, .31] | .64, [.63, .65] | .6, [.59, .61] |
| ECG | .52, [.5, .54] | .18, [.18, .19] | .27, [.27, .28] | .59, [.58, .6] | .57, [.56, .58] |
| EYE | .61, [.59, .64] | .16, [.16, .16] | .25, [.25, .26] | .56, [.55, .57] | .54, [.53, .55] |
| EDA | .52, [.5, .54] | .16, [.16, .17] | .25, [.24, .26] | .56, [.55, .57] | .54, [.53, .55] |
| RSP | .6, [.58, .62] | .15, [.14, .15] | .24, [.23, .24] | .52, [.51, .53] | .51, [.51, .52] |

Supplementary Discussion on Methodology

Our analysis examined the whole 110s period preceding a mental state report. The rationale for this epoch choice was that specific features from slow oscillatory body afferents, such as heart rate variability, cannot be reliably estimated from shorter time windows. Our results advance the discussion of how long mental states last. The study of a mental state is based on a pre-probe window preceding a mental report, akin to our paradigm. The examined pre-probe epoch is varied based on the neuronal or electrophysiological modality used, as BOLD markers are typically slower than EEG. This approach has shown that mental states can be decoded as early as 10s before report [8, 9, 10], or as late as 500ms before the report [11]. Here we show that information about a mental state might be available even earlier. However, an important caveat to account for this interpretation is that experience-sampling introduces a discrete methodology to sample mental states. Therefore, our analysis of the 110s might encompass multiple mental states, and the patterns of brain-body might represent those as well. To account for this, we performed exploratory analysis by classifying MB considering only a pre-probe period of 10s. The new classifier did not show improved performance. Instead, we observed comparable results across the 10s and the 110s classifier when pooling data across BRAIN and BODY markers. Similarly to the 110s classifier, we examined whether specific BRAIN or BODY markers could decode MB reports separately from other domains of features. While EEG retained high performance, we observed a significant drop in performance across all BODY classifiers, and most prominently, in the eye classifier. Therefore, we speculate that a short preprobe period might not be adequate to reliably estimate informative BODY features for MB classification.

Pooling together MB reports across different arousal conditions might hide arousal-specific effects on MB reports. To examine this hypothesis, we performed exploratory analyses by training our classifier separately on different arousal conditions. While we were able to retain predictive power across these partitions, we observed distinct different brain-body feature importance. During baseline, the feature importance ranking was comprised of a diverse combination spanning EEG, ECG, and EYE markers. There was a clear shift in importance in sleep deprivation, with the power of the delta band being the most important feature. This result is consistent with contemporary accounts of MB [11, 12], where MB is correlated with the presence of the presence of parietal slow-wave intrusions in wakefulness, within the delta band. Mounting sleep pressure has been correlated with an overall increase in localized sleep-like slow waves [13, 14]. As these intrusions in the delta band increase with sleep deprivation, attentional lapses in the form of MB might become themselves more intrusive to cognition, and account for the overall cognitive decline in performance during sleep deprivation [15]. Contrary to sleep deprivation, post-exercise MB has significantly different feature importance, not driven by delta, but by connectivity in alpha and theta bands, as well as contributions for EDA and EYE markers. Reductions in alpha and theta bands are consistent with reduced arousal after propofol-induced anesthesia [16], consistent with the hypothesis of altered arousal during MB. Meanwhile, contributions of the phasic component of the EDA signal provide direct evidence for sympathetic contributions to MB.
